# CIVET-Macaque: an automated pipeline for MRI-based cortical surface generation and cortical thickness in macaques

**DOI:** 10.1101/2020.08.04.237149

**Authors:** Claude Lepage, Konrad Wagstyl, Benjamin Jung, Jakob Seidlitz, Caleb Sponheim, Leslie Ungerleider, Xindi Wang, Alan C. Evans, Adam Messinger

## Abstract

The MNI CIVET pipeline for automated extraction of cortical surfaces and evaluation of cortical thickness from *in-vivo* human MRI has been extended for processing macaque brains. Processing is performed based on the NIMH Macaque Template (NMT), as the reference template, with the anatomical parcellation of the surface following the D99 and CHARM atlases. The modifications needed to adapt CIVET to the macaque brain are detailed. Results have been obtained using CIVET-macaque to process the anatomical scans of the 31 macaques used to generate the NMT and another 95 macaques from the PRIME-DE initiative. It is anticipated that the open usage of CIVET-macaque will promote collaborative efforts in data collection and processing, sharing, and automated analyses from which the non-human primate brain imaging field will advance.

## 1. Introduction

The cerebral cortex has a two-dimensional folded topology, tangentially subdivided into cortical areas and radially into layers. Human neuroimaging studies of cortical structure and function have been advanced through the development of automated surface reconstruction techniques (Kim et al. 2005; Bruce Fischl 2012). These techniques have enabled intersubject and intermodal coregistration, compression and analysis of large datasets, and revealed areal and laminar organization in a common surface-based coordinate system.

Cortical meshes enable quantification of morphometry: per-vertex cortical thickness, curvature, and surface area can all be readily measured from mesh representations. The spherical topology of the cortical surfaces allow for intersubject coregistration (B. Fischl et al. 1999; Lerch and Evans 2005; Robinson et al. 2014). Surfaces can be aligned to a common average template, enabling group-comparison studies, and surface-based parcellations such as atlases can be mapped to individual surface meshes for regional analyses, with per-vertex correspondence between the white matter and pial surfaces. Furthermore, coregistering other imaging modalities to the same space enables quantitative sampling of any number of *in vivo* modalities: fMRI, qMRI, PET, EEG, MEG (Weiskopf et al. 2013; Greve et al. 2014; Bonaiuto et al. 2018; Yeo et al. 2011), as well as *post mortem* histological (Wagstyl et al. 2020) and genetic data (Hawrylycz et al. 2012). Once sampled to the mesh, and coregistered to a common template, vertex-based representations allow for intermodal analysis and application of more complex analytical techniques.

Such techniques have been used to advance human neuroimaging, and have the potential to similarly advance NHP neuroimaging. For instance, in humans, surface analysis has been used to parcellate the cortex according to multimodal *in vivo* data from task and resting state fMRI and structural images in the same subjects (Glasser et al. 2016). Radial sampling between gray and white cortical surfaces across the cortex can produce depth-dependent intensity profiles that, with sufficient resolution and contrast, can be used to segment cortical layers (Wagstyl et al. 2020). Surface-based descriptions have also been used to characterize human semantic organization (Huth et al. 2016) and pathology (Lerch et al. 2005). In NHP studies, surfaces can be used to identify similar regions across species and facilitate interspecies comparison (Xu et al. 2019; Miranda-Dominguez et al. 2014; Donahue et al. 2018). Thus, cortical surface reconstruction techniques provide a powerful framework for neuroimaging.

Several cortical surface reconstructions for macaques have been presented (Koo et al. 2012; Oguz et al. 2015; Donahue et al. 2018; Wang et al. 2018; Autio et al. 2020) and a few new pipelines for macaque cortical surface generation have recently been shared on the PRIME-Resource Exchange (https://prime-re.github.io/) (Messinger et al., *this issue*). These efforts have made surface generation less labor intensive. A surface-free approach to evaluate cortical thickness, using the DiReCT volume registration-based method in ANTS (Tustison et al. 2014), has also been extended to the macaque (Calabrese et al. 2015). However, most of the procedures described require both T1 and T2-weighted anatomical scans of sufficient quality, multiple software packages, various manual interventions, and several end-arounds to get human software tools to process non-human primate data. Furthermore, these packages are limited to exemplary scans or small datasets, some without means for surface-based registration for group analyses of large cohorts. The work presented here addresses the need for an open-access dedicated pipeline for non-human primate cortical surface reconstruction, with capabilities comparable to human pipelines.

The CIVET pipeline (Lepage et al. 2017), which was developed at the Montreal Neurological Institute (MNI), is a comprehensive automated pipeline for corticometric and morphometric analyses of human MR images. The pipeline provides high resolution cortical surface reconstruction from T1-weighted MR images, featuring intersubject surface-based coregistration with per-vertex correspondence between gray and white matter surfaces. In this work, CIVET has been customized for the processing of non-human primate MR images. The present work covers the extension to the macaque monkey and is based on the NIMH Macaque Template (NMT) (Seidlitz et al. 2018).

The availability of a new automated pipeline for processing non-human primate anatomical scans opens new opportunities for analyzing large datasets with their wealth of structural, functional, diffusion and post mortem data. For example,the UNC-Wisconsin Rhesus Macaque Neurodevelopmental Database provides a rich dataset of longitudinal structural and diffusion MRI during early postnatal development in 34 rhesus macaques (Young et al. 2017). For post mortem data, the Allen Brain Institute provides gene expression and histological data throughout macaque developmental stages, with reference MRI (Bakken et al. 2016). Others have published postmortem diffusion tensor imaging dataset (Calabrese et al. 2015). Recent initiatives such as the PRIMatE Data Exchange (PRIME-DE) (Milham et al. 2018) have helped to harmonize these data, generating large scale neuroimaging datasets that require robust, automated analytical pipelines. Furthermore, adaptation of statistical map repositories like Neurovault (Gorgolewski et al. 2015) have made it easier to share standardized volumetric results in NHPs. Analysis of such data in terms of anatomical areas and morphometry would be enhanced by surface analysis capabilities (Calabrese et al. 2015). Even existing surface-based studies of NHPs may benefit from improved techniques and automation (Koo et al. 2012).

## 2. Material and Methods

### 2.1 Subject Information

This study uses anatomical MR scans from 3 sources: the *ex-vivo* scan of the single-subject D99 template (Reveley et al. 2017), 31 macaques from the Central Animal Facility at the National Institute of Mental Health (NIMH, USA), and *in-vivo* T1-weighted (T1w) anatomical scans of 95 macaques obtained from the INDI PRIME-DE consortium (Milham et al. 2018, http://fcon_1000.projects.nitrc.org/indi/indiPRIME.html). All animal procedures were conducted in compliance with the National Institutes of Health Guide for the Care and Use of Laboratory Animals or, in the case of the PRIME-DE scans, with the animal care and use policies of the institution where the data was collected (Table 1).

**Table 1:** Scanning information for 95 selected T1w scans from the INDI PRIME-DE consortium. Only scans obtained in the sphinx position with complete published information about age and sex were retained.

| Site | N (M/F) | Age range (yr) | Scanner Type |
| --- | --- | --- | --- |
| 1-Newcastle University | 12 (10/2) | 5-13 | Bruker Vertical, 4.7T |
| 2-NIMH (Leopold, Russ) | 2 (1/1) | 6-7 | Bruker Vertical, 4.7T |
| 3-Princeton University | 2 (2/0) | 3 | Siemens Prisma, 3T |
| 4-University of Minnesota | 2 (0/2) | 10+ | Siemens Syngo, 7T |
| 5-UC Davis | 19 (0/19) | 18-22 | Siemens Skyra, 3T |
| 6-NKI | 2 (1/1) | 6-7 | Siemens TIM Trio, 3T |
| 7-Mount Sinai | 9 (8/1) | 3-8 | Philips Achieva, 3T |
| 8-Mount Sinai | 5 (5/0) | 5-6 | Siemens Skyra, 3T |
| 9-Oxford University | 20 (20/0) | 2-6 | n/a, 3T |
| 10-Western Ontario | 3 (3/0) | 4-7 | Siemens Magnetom, 7T |
| 11-Institute of Neuroscience | 8 (7/1) | 3-5 | Siemens TIM Trio, 3T |
| 12-McGill University | 1 (0/1) | 12 | Siemens TIM Trio, 3T |
| 13-East China Normal University-Kwok | 4 (4/0) | 2-3 | Siemens TIM Trio, 3T |
| 14-Aix-Marseille Universite | 4 (3/1) | 7-8 | Siemens Prisma, 3T |
| 15-Netherlands Institute for Neuroscience | 2 (2/0) | 5 | Philips Ingenia, 3T |

The rhesus macaques (*Macaca mulatta*) scans from the NIMH, referred to hereafter as the NMT cohort, have been previously described (Seidlitz et al. 2018). The 25 male and 6 female animals were, at the time of the scans, juveniles and adults (5.2 ± 1.9 years) with an average weight of 6.3 ± 1.7 kg. The animals did not have surgery or procedures within the intracranial cavity prior to the scans. However, 14 had a non-metallic post surgically implanted on top of their skull that made the dorsal portions of the skull appear thin or absent in some scans. T1w MR volumes were acquired using the modified driven equilibrium Fourier transform (MDEFT) sequence (Lee et al. 1995; Deichmann, Schwarzbauer, and Turner 2004) on a 4.7T horizontal scanner (Bruker Biospec 47/40). On the day of scanning, animals were placed under food restriction, anesthetized with isoflurane, and placed in the scanner in the sphinx position. Each whole-brain MDEFT scan was acquired over a 40-60 minute session using either a 14 or 16.5 cm diameter single loop circular head coil (FOV = 96 × 96 × 70 mm^3^; 0.5 mm isotropic voxels). For most subjects, multiple MDEFT scans were collected consecutively during the session (mean = 2.5 MDEFT scans/subject, range = 1-7). In these cases, subsequent MDEFT scans were linearly (6 parameter rigid-body) registered to the first, and then averaged to create a single volume for each subject.

The scans for the INDI PRIME-DE consortium consisted of *Macaca mulatta* (N=90) and *macaca fascicularis* (N=5) monkeys and were collected at various sites with different scanner types and field strengths. All scans retained for this study (T1w only) were collected in the sphinx position and had published age and sex information (see Table 1). Scans of very poor quality or with obvious lesions were excluded from this experiment.

### 2.2 Overview

The CIVET pipeline for extraction of cortical surfaces from in-vivo MR images of the human brain has been extended for processing macaque brains, and as such the processing stages for macaques parallel those for the human. Extensions of CIVET to the macaque use the NIMH Macaque Template (NMT) as the standardized space for extracting and characterizing the surfaces of individual monkeys (Seidlitz et al. 2018). The macaque option for CIVET, herein referred to as CIVET-macaque, is invoked by the simple specification of a command line parameter for the choice of the model for the standardized space. CIVET takes as input a T1w scan and registers if to standardized space, herein the NMT (Seidlitz et al. 2018), using an affine transformation. The image is resampled into the NMT space, corrected for non-uniformities, masked, then classified into cerebrospinal fluid (CSF), cortical and subcortical gray matter (GM), and white matter (WM). A white matter surface, per hemisphere, is fitted at the GM-WM boundary using a marching-cubes algorithm, then adjusted to the maximum T1w intensity gradient position, before being expanded to the pial surface. Cortical thickness is calculated as the distance between the two cortical surfaces. Surface-based registration is performed and the cortical thickness maps are blurred and resampled at the common vertex locations of the template for group comparisons and analyses.

The pipeline is based on the *minc* storage format developed at the MNI (Vincent et al. 2016) as the processing unit. The *minc-2* format is based on the HDF5 library with automatic internal file compression for efficient data representation of large volumes (The HDF Group 2016). Cortical surfaces are represented using the *MNI-obj* format (http://www.bic.mni.mcgill.ca/Services/HowToWorkWithObjectFiles). Many tools to manipulate and visualize volumes and surfaces are available in the openly available *minc toolkit* (http://bic-mni.github.io/#MINC-Toolkit). An option has also been added to CIVET to export its outputs to the commonly used NIfTI/GIfTI formats, enabling compatibility with a wide range of other analysis tools. The CIVET pipeline is available on the CBRAIN platform (Sherif et al. 2014) for collaborative studies and sources and binaries for Linux platforms are available via the CIVET project page (https://github.com/aces/CIVET_Full_Project).

While the processing flow in CIVET is the same for humans and macaques, the main implementation issues in the extension of CIVET to macaques are the elaboration of species-specific volumetric and surface templates for registration as well as the scaling of all length-related parameters. Numerous modules of the pipeline have been modified to include a length scale parameter to account for the smaller brain size in the macaque. A linear factor of 0.4 (relative to the human brain) is used for the macaque. Length scales appear in image blurring (full width at half maximum, fwhm), registration (search distances), masking (search radius), surface blurring of cortical thickness (fwhm), morphological operations such as voxel erosion and dilation (distance), non-uniformity corrections (spline distance), and voxel size representation proportional to brain structures. Other algorithms such as tissue classification (histogram-based) and cortical surface extraction (topology based) are unaffected. These changes are summarized in Table 2. Further adaptations of the pipeline to other non-human primate brains should be straightforward by introducing the appropriate length scales and supplying species specific templates.

**Table 2:** Differences in processing steps between the CIVET-human and CIVET-macaque configurations.

| Processing Step | Human Processing | Macaque Processing |
| --- | --- | --- |
| Stereotaxic template | MNI-ICBM, ADNI, Colin27 | NMT |
| Stereotaxic voxel sampling | 0.50mm or 1.0mm | 0.25mm or 0.50mm |
| Volumetric registration (fwhm, simplex, sampling) | length scale 1.0 | length scale 0.4 |
| Neck cropping (headheight) | 175mm | 50mm |
| Masking (search distance) | length scale 1.0 | length scale 0.4 |
| Non-uniformity corrections | spline distance 125 (3T) | spline distance 50 (4.7T) |
| Classification | priors on human template | priors on NMT |
| Segmented masks (brainstem, ventricles, cerebellum, etc.) | on human template | on NMT |
| Surface extraction (fwhm, gradient, dilation/erosion) | length scale 1.0 | length scale 0.4 |
| Surface resolution | 40,962 or 163,842 vertices | 40,962 or 163,842 vertices |
| Cortical thickness method | tlink, tlaplace, tfs | tlink, tlaplace, tfs |
| Cortical thickness blurring | 30mm fwhm | 8mm fwhm |
| Surface registration (fwhm) | length scale 1.0 | length scale 0.4 |
| Surface parcellation | AAL, DKT, lobes | CHARM/D99, lobes |

### 2.3 Volumetric template

The first step in the extension of CIVET is the selection of a volumetric template for stereotaxic registration and the preparation of its associated atlases. The NIMH Macaque Template (NMT) version 1.2 (asymmetric model), built from the non-linear average of the 31 scans from the NMT cohort, was selected as this registration target (https://afni.nimh.nih.gov/NMT; Seidlitz et al. 2018). We note here that, while it is not used in CIVET, the NMT has been recently updated and now includes a symmetric macaque template in Horsley-Clarke stereotaxic coordinates (Horsley and Clarke 1908; Jung et al., *this issue*). The NMT (version 1.2) was created by iterative symmetric diffeomorphic registration methods using the ANTs software package (Avants et al. 2010). First, whole-head single subject images were initially aligned to a standard coordinate space (D99-SL from (Reveley et al. 2017)), and resampled from 0.5 to 0.25 mm isotropic resolution. Then, following a voxelwise N4 bias field correction (Avants et al. 2011) applied on each subject’s aligned image, a spatially-unbiased population average was created using the symmetric group-wise normalization (SyGN; (Avants et al. 2010; Love et al. 2016)). The resultant NMT was placed in the neurological orientation, rotated so that, like the D99 atlas, the horizontal plane contained both the superior extent of the anterior commissure and the inferior extent of the posterior commissure (Horsley and Clarke 1908; Saleem and Logothetis 2012), and translated to place its origin (AP 0, SI 0, ML 0) on the midline at the center of the anterior commissure. Intensity values in the NMT were capped to account for the extreme values of the blood vessels, and a final N4 bias field correction was performed.

Atlases on the NMT (version 1.2) are created for its use in CIVET. The template is segmented to label the cerebellum, brainstem, ventricles, amygdala, hippocampus, and subcortical region.

This segmentation is used in CIVET to exclude the brainstem and cerebellum during tissue classification and to mask out the ventricles and subcortical gray areas prior to the extraction of the white matter surface. A brain mask of the template is obtained using a tailored version of FSL *bet* for macaques (Smith 2002). Finally, tissue priors are defined on the NMT (500+ points for each of CSF, GM, and WM). Priors are also defined in the background (BG) outside the skull. These priors are later coregistered to each subject and used for training the tissue classifier. The pre-processing steps of the template are summarized in Figure 1.

**Figure 1:**
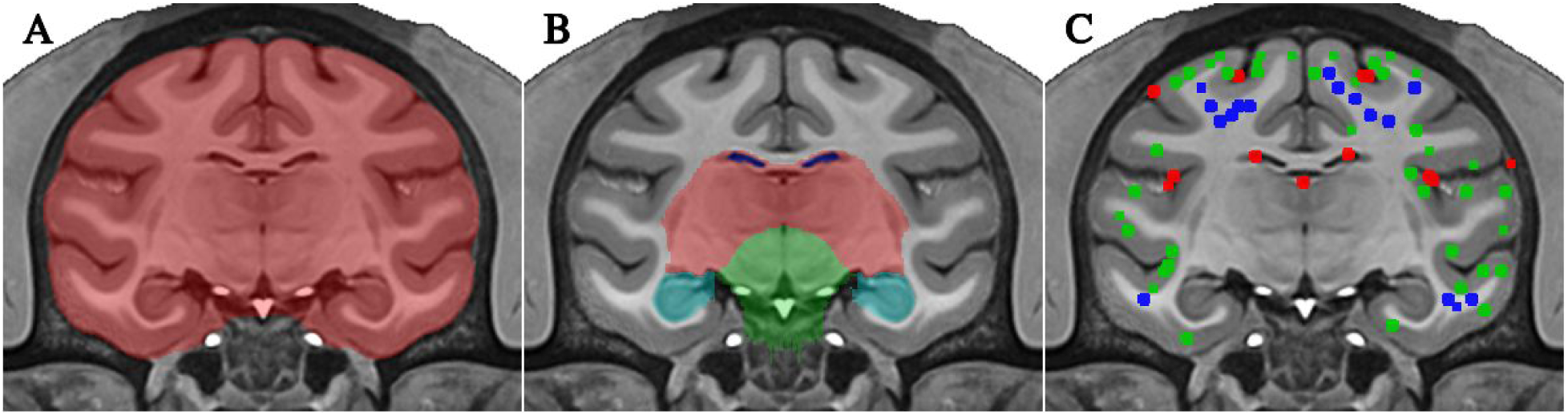
Pre-processing of NMT: A) brain mask (red); B) regional masks for ventricles (dark blue), subcortical region (red), brainstem (green), cerebellum (not visible in this plane), hippocampus (cyan), and amygdala (not visible in this plane); C) subset of the points used to define priors for the three tissue classes: CSF (red), GM (green), WM (blue).

### 2.4 Registration to standardized template space

Registration to the NMT uses *minctracc* (Collins et al. 1994). *Linear registration follows a hierarchical multi-scale approach from coarse to fine image blurring and sampling rates. For the macaque, all length scales in the registration procedure are scaled by 0.4 to account for brain size relative to that of the human. That is, the fwhm* for image blurring and sampling intervals are scaled proportionally by 0.4. With this scaling, the registration procedure for the human brain achieves optimal performance on the macaque brain. The length scales in the non-linear registration procedure are adjusted in the same way.

Linear and non-linear registration to the macaque template requires the use of a brain mask since head size varies significantly between males and females, unlike humans, resulting in variable scan field-of-views. For instance, the male macaque typically has larger temporalis muscles than the female, resulting in thicker “scalp” (muscle) and head size. In some cases, brains have been pre-processed and de-skulled. Registration, using *minctracc*, proceeds by specifying the brain mask on the target image only, herein the NMT volume with its associated brain mask, and evaluating the source image, herein the subject’s scan, at the sampling points within the target mask, without the need to have a brain mask explicitly defined on the subject. This registration approach has proven to be robust in overcoming the variability in scans due to head size and field-of-view. In the rare case when registration fails, a custom brain mask in native space can be supplied. This custom brain mask can be obtained from automated tools within the CIVET environment or by other external tools. The accuracy of such custom masks should be assessed by the user prior to processing.

The macaque brain is resampled in linear standardized space at the processing voxel resolution of 0.25mm or 0.50mm, which is of comparable resolution relative to the human brain (processing resolution at 0.50mm or 1.0mm).

### 2.5 Non-uniformity correction

The image, resampled into standardized space, is corrected for non-uniformities using N3 bias field correction (Sled, Zijdenbos, and Evans 1998). The sampling region for N3 is restricted by the brain mask of the volumetric template, as it is for humans, since a reliable custom mask cannot be obtained prior to N3 correction. The parametric spline distance must be entered by the user. This distance varies with the field strength of the scanner and, to some extent, the species’ head size. Finally, denoising is performed on the input image to smooth out voxels with local maxima of intensity (100 iterations), thereby avoiding the inclusion of blood vessels in the calculation of the bias field.

### 2.6 Brain-masking

Brain-masking is based on extensions of FSL *bet* version 1 (Smith 2002). Given the differences in total head size between macaque males and females, it is critical to exclude the scalp prior to calculating the histogram of intensities for establishing the masking thresholds. The most reliable approach was to sample the intensities within the brain mask of the volumetric template, which is permissible in standardized space. Brain extraction a la FSL *bet* thus provides a refinement of the template’s brain mask. For some scanning protocols, the blood vessels have very large signal intensities that interfere with convergence and result in an incomplete mask. To prevent this, denoising is performed on the image to smooth out voxels with local maxima of intensity (100 iterations). Moreover, the brain-masking procedure can incorporate information from complementary MR scans, such as a T2-weighted image, for improving the T1w brain mask. The procedure proved to be very robust at extracting accurate masks on scans of individual subjects (see Figure 2).

**Figure 2:**
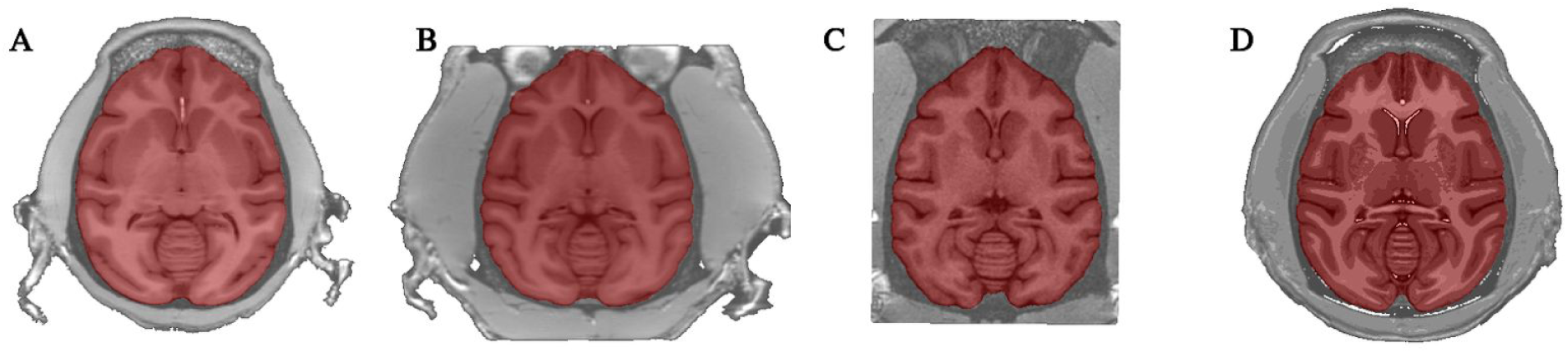
Automated extraction of brain mask in NMT standardized space on sample scans of the NMT cohort, highlighting head size differences in male and female macaques and inter-scan field-of-view variability: A) female macaque; B) male macaque; C) male macaque with cropped field of view; and D) the NMT volume itself (25 males, 6 females). All masks are perfectly extracted regardless of the field of view of the original scan and of the head size of the animal.

### 2.7 Tissue classification

The tissue classification procedure is identical for both human and macaque scans and relies on the T1w scan, with the possibility of a multi-modal classification should T2-weighted and/or proton density scans be available. The brain is classified into cerebrospinal fluid, gray matter, and white matter using an artificial neural network (ANN) trained on the intensities of the tissue priors (BG, CSF, GM, WM) mapped non-linearly from the NMT volume to the subject (Zijdenbos, Forghani, and Evans 1998). The subcortical region labelled on the NMT allows for the discrimination of cortical from non-cortical gray matter in the specific region. Partial volume estimates are obtained for the tissue classes (Tohka, Zijdenbos, and Evans 2004). Partial volumes for CSF are combined with curvatures (eigenvalues of the Hessian matrix of intensities) in the reconstruction of the CSF skeleton delineating buried sulci which ensures full sulcal penetration of the pial surface during its creation.

### 2.8 Masking blood vessels

Prior to the extraction of the white matter surface, it is critical to identify T1w hyperintensities, commonly representing major blood vessels and erroneously segmented as white matter by the classifier, to prevent these spurious white matter voxels from interfering with the process of extracting the white matter surface. The basic assumptions in the detection algorithm are that blood vessels are represented by T1w hyperintensities with a line morphology propagating in CSF within sulci. Sulcal lines are identified using a watershed algorithm in non-white voxels, then dilated partly through gray matter and, consequently, through blood vessels, but not white matter. The trace of the Hessian matrix of second derivatives (i.e., the sum of the Hessian’s three eigenvalues) is used as an approximation of the local curvatures to detect one-dimensional line features. Thresholds for the expected signal intensity in white matter and for the trace of the Hessian are used to discriminate lines structures in sulci from blood vessels. Figure 3 illustrates blood vessels detected on a single scan.

**Figure 3:**
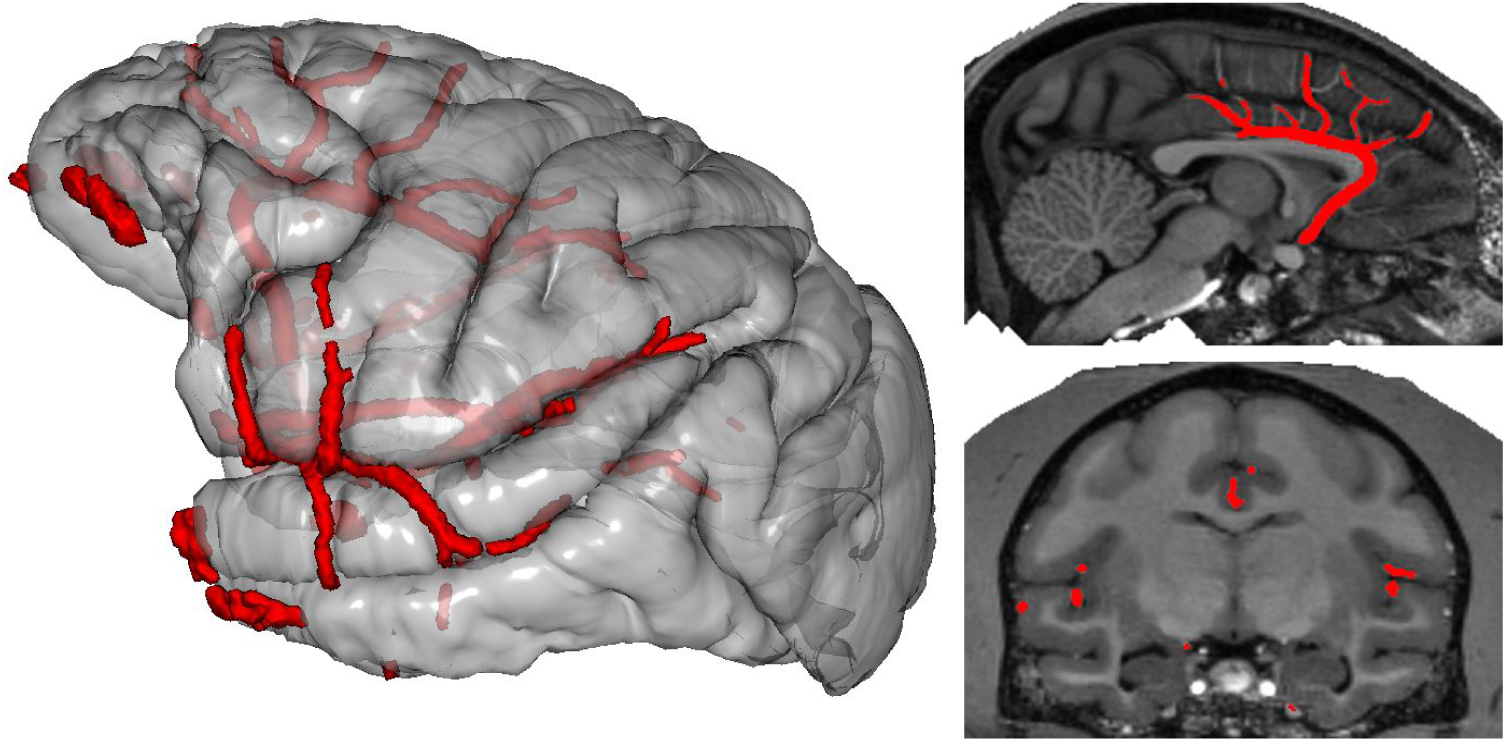
Major blood vessels identified from T1w hyperintensities for one NMT subject. Left: representation of blood vessels (red) on a semi-transparent lateral view of the pial surface. Right: T1w hyperintensities revealing the blood vessels in a sagittal and coronal slice through the volume. The vessel-like hyperintensities are discounted during extraction of the white matter surface.

### 2.9 Cortical surface extraction

A white matter mask is created from the tissue classification, masked for the brain. Using the segmented regional masks on the NMT volume non-linearly transformed to the subject, the ventricles and subcortical gray matter are filled in as white matter, the hippocampus and the amygdala are masked out, and the brainstem and cerebellum are excluded for the purpose of extracting the white matter surface. The brain, in NMT space, is then split into the left and right hemispheres, at the mid-plane wall, and the white matter surfaces are extracted using a marching-cubes algorithm (Lepage et al. 2017). Spherical topology of the cortical mantle is preserved by advancing the surface, in voxel space, outside-in from a sphere enclosing the hemisphere, hence the requirement for masking off the blood vessels obstructing the progression of the surface. Mesh adaptation operations (vertex smoothing, edge collapsing and swapping) are used to efficiently reduce the surface mesh to a manageable size (around 140,000 vertices) for further processing. The decimated white matter surface is inflated to a sphere and resampled at a fixed number of vertices on a canonical sphere obtained by refinement of the icosahedron (40,962 vertices). The resampled surface is then registered to the average white matter surface template (see subsection below) for mapping the surface region on which the intensity gradient correction is disabled, either because it is meaningless (cut through the corpus callosum and brainstem on the medial wall) or misleading (e.g., cavity of the hippocampus and amygdala, insula region with low gray-white contrast in proximity of the claustrum). The white matter surface is then adjusted to the maximum intensity gradient (magnitude of first derivative of T1w intensity normal to the surface).

The pial surface is obtained by expanding the white matter surface radially outwards up to the pial boundary using the CLASP algorithm (Kim et al. 2005). Consequently, a per-vertex correspondence exists linking the vertices of the white matter and pial surfaces, from which the mid-surface is obtained by averaging the position of the linked vertices. Finally, during the construction process, the right surface is obtained by mirroring the canonical sphere before resampling, thus establishing a direct correspondence between vertices across hemispheres at mirror-symmetric positions.

No special adjustments were made for macaque surface extraction except for scaling (by 0.4) the search distance for the gradient correction and the *fwhm* for blurring volumes. The surfaces can also be extracted at high-resolution (163,842 vertices).

### 2.10 Surface-based registration

Surface-based registration is used to find the correspondence between the cortical surface of an individual subject to a surface template (Robbins 2003). The registration operates on a spherical representation of the surfaces and the mesh alignment is driven by a *depth potential* term, akin to sulcal depth (Boucher, Whitesides, and Evans 2009). Thus sulci are aligned to sulci and gyri are aligned to gyri. Following registration, any map defined on the subject’s surface can be interpolated at the sampling vertices of the reference surface template. Resampling of surface maps, for instance cortical thickness, enables vertex-wise group comparisons across subjects of a population. Furthermore, regional analyses can be performed based on parcellations defined on the template. The generalization from human to macaque required accounting for head size by scaling the fwhm for blurring the sulcal depth term on the surface.

Surface-based registration uses a white matter surface template for the gradient correction adjustment of the white matter surface and a mid-surface template for resampling surface maps such as cortical thickness, curvature, etc. The next subsections describe the creation of these average white and mid-surface templates as the targets for surface-based registration and the application of the D99 and CHARM parcellation on these surface templates.

### 2.11 Average surface template

The average surface template is generated from the NMT dataset (N=29, 2 subjects were rejected -- one with a lesion, the second one due to abnormal ventricles) as the target for surface-based registration. Like for the human brain, the average surface is constructed by averaging the surfaces of the individual subjects, as opposed to using the cortical surfaces extracted on the average volume of all subjects. But, unlike the human brain, the average surface is generated in the non-linear standardized space, since subject-to-subject sulcal variability is low for the macaque. That is, the non-linear transformation from the subject’s volume to the NMT template is applied to the subject’s surfaces before averaging them, thus reducing geometrical variance. This low-frequency non-linear transformation essentially corrects for gross distortion, like global twist, while retaining micro shape differences describing biological variability (see Figure 4). Non-linear warping was also applied in the construction of the NMT volume (Seidlitz et al. 2018).

**Figure 4:**
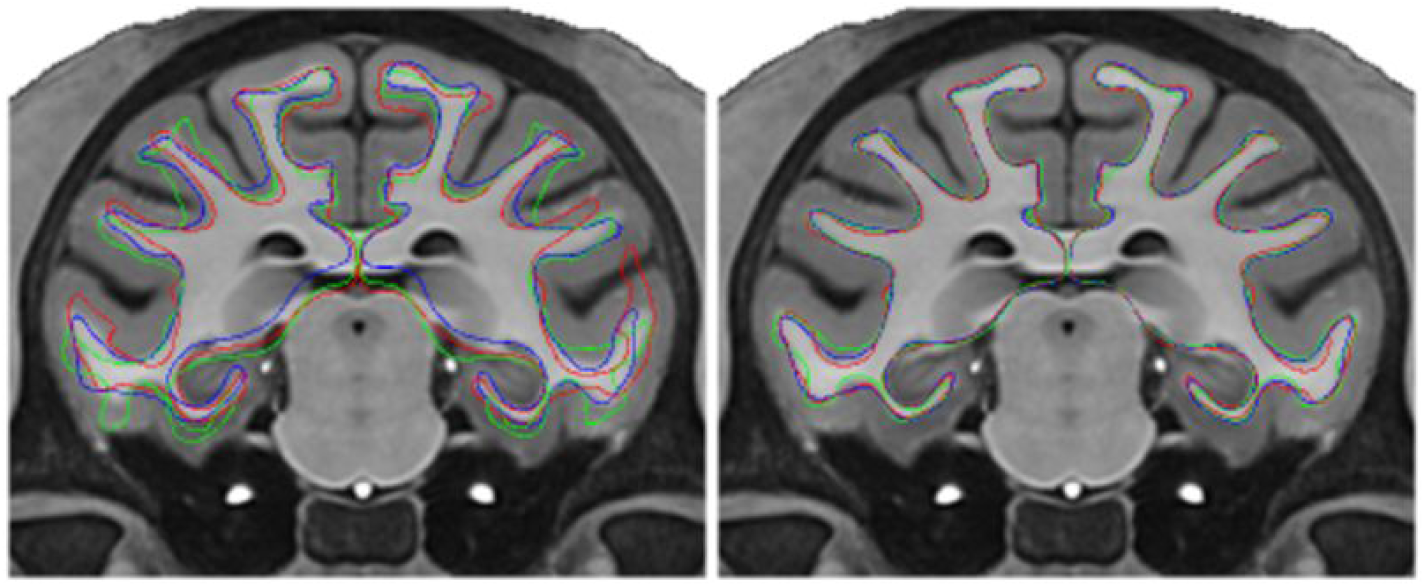
View of white matter surfaces for 3 NMT subjects in linear (affine) space on the left and in non-linear space on the right. Averaging in non-linear space eliminates shape differences while retaining the full cortical morphology of the macaque brain during the creation of the average white matter surface. The surfaces are overlaid onto the T1w NMT. The colored lines correspond to the three subjects.

The creation of the average surface is cyclic: CIVET requires the existence of an average surface as a template for surface-based registration, but surfaces must exist prior to averaging them. An iterative process is followed by first averaging unregistered surfaces and then using this average as the target for registration on the next iteration, and so on. The initial average is made possible since all surfaces have the same number of vertices, via sphere-to-sphere interpolation of the inflated surface to a canonical sphere, and that coordinates are aligned in stereotaxic space (Lyttelton et al. 2007).

The generation of the average white matter surface using the marching-cubes algorithm is computationally intensive since the algorithm requires repeated processing steps at each averaging iteration, namely, extraction, mesh decimation, inflation to sphere, resampling, and surface-based registration in order to apply the mask for the gradient correction. This mask must be defined on an intermediate average surface used as the registration target, which itself evolves as the average surface is iteratively converged. As such, the convergence of the individual white matter surfaces depends on the average surface at each iteration and these white matter surfaces must therefore be recomputed in full at each iteration, making the overall process lengthy. To accelerate the convergence process, the intensity gradient correction is disabled for the first 5 iterations of the creation of the average white matter surface. The gradient mask is defined on the intermediate average white matter surface and frozen through 5 more global iterations of surface averaging, with the intensity gradient correction enabled. Full convergence is rapidly achieved (see Figure 5).

**Figure 5:**
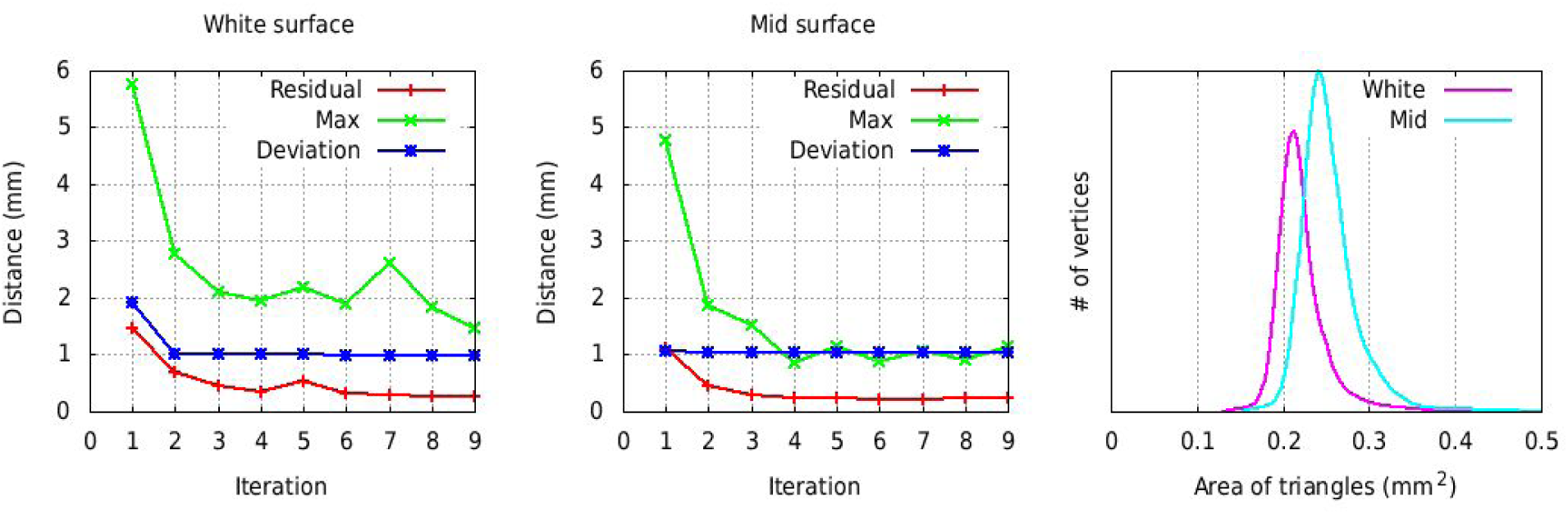
Convergence of the average symmetric white matter (left panel) and mid-cortical (middle panel) surfaces for the NMT cohort (N=29). The residual represents the vertex-wise mean displacement in the average surface from one iteration to the next and should tend to zero as the vertices are no longer moving. Max represents the largest vertex-wise displacement from one iteration to the next. Deviation represents the mean vertex-wise displacement for the N=29 subjects (58 surfaces) and is a measurement of inter-subject shape variability. The rightmost panel shows the histogram of the area of the triangles on the final average white matter and mid-cortical surfaces, indicating a nearly uniform point sampling over the surface.

The creation of the average mid-surface is much simplified since the extraction of the cortical surfaces does not need to be repeated at each iteration. On start-up, the mid-surfaces are initially registered to the average white matter surface to align the vertices between the two surfaces. Ten global iterations are used in the process, but convergence is achieved after only a few iterations (see Figure 5).

For the symmetric averages, the right surfaces are mirrored as left surfaces before being averaged. For both white matter and mid average surfaces, the vertices are equidistributed on the surface to obtain a surface with nearly equal-size triangles, with the goal to have uniform sampling over the entire cortical sheet (see Figure 5). The area equalization process is performed at the end of each averaging iteration and consists of moving each vertex to the centroid of its neighboring vertices, weighted by the area of their connected triangles.

The average white matter surface is shown in Figure 6. The variability in shape (mean vertex displacement) across the N=29 subjects is highest in the parieto-occipital area and the medial and ventral occipital lobe. The average mid-cortical surface exhibits the same properties as the average white matter surface.

**Figure 6:**
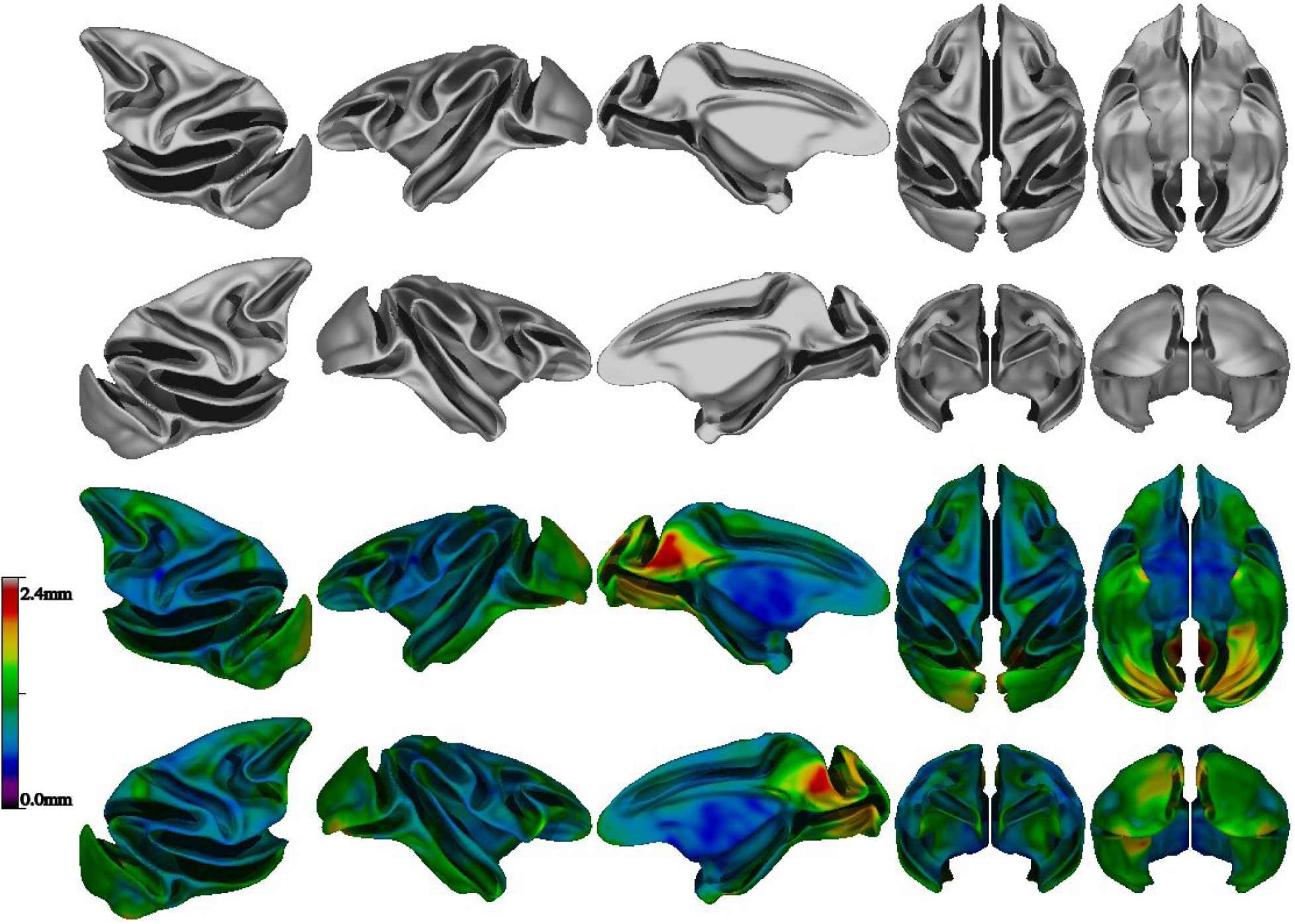
View of the average white symmetric surface colored by mean vertex displacement for N=29 subjects of the NMT dataset. Higher variability in shape was observed in the parieto-occipital area (visual areas V6/V6A) as well as in the occipital lobes, areas of substantial inter-subject differences in macaques.

### 2.12 Cortical morphometry

The cortical surfaces, extracted in standardized space, are transformed back to native space and cortical thickness is evaluated as the distance between the white matter and pial surfaces. Three cortical thickness metrics are available in CIVET:

1. *tlink*: distance between linked vertices between the white matter and pial surfaces (sensitive to mesh distortion during expansion of the pial surface, which gives rise to an overestimation of cortical thickness);
2. *tlaplace*: distance along radial lines defined by a Laplacian field between the white matter and pial surfaces (not sensitive to mesh distortion) (Lerch and Evans 2005);
3. *tfs*: symmetric average of projected distance from white matter-to-pial and pial-to-white matter surfaces (like FreeSurfer).

Other metrics are also produced such as vertex-wise surface areas and volumes, and mean and Gaussian curvatures. These measurements are blurred isotropically on the mid-cortical surface in native space (Boucher, Whitesides, and Evans 2009). Default values for surface blurring (*fwhm*) are proportionally less for the macaque than for the human (e.g. fwhm of 8mm). All measurements are resampled on the average NMT symmetric surface template as a basis for group comparison analyses.

### 2.13 CHARM surface atlas

The D99 subject was processed with CIVET-macaque to obtain the D99 surface parcellation (Reveley et al. 2017). This post-mortem brain scan has no skull, a T1w-like contrast, and has been pre-processed: non-uniformities have been corrected and the scan has been symmetrized and transformed to a standardized space. A value of 0 was used as the spline parametric distance to disable N3 and the blood-vessel algorithm has been disabled as well (because the scan was acquired *ex vivo*). Surface-based registration was used to register the D99 mid-surfaces to the NMT symmetric surface template to ensure a direct vertex-to-vertex correspondence. The resampled D99 mid-cortical surfaces were then intersected with the D99 volumetric segmentation (Reveley et al. 2017) to obtain parcellation labels, which mapped directly to the NMT symmetric surface template (see Figure 7). A few minor morphological operations on the extracted labels were necessary to enforce symmetry and to correct for mis-labeled and unassigned vertices arising when intersecting the surfaces with the labelled volume. Using the final D99 labels, each vertex was assigned 6 additional anatomical labels; one for each level of the Cortical Hierarchical Atlas of the Rhesus Macaque (CHARM) (Jung et al., *this issue*). The 6 anatomical levels of CHARM range from a lobar parcellation at the coarsest level (4 ROIs) to nearly the full set of D99 cortical areas at the finest level (139 ROIs). Figure 8 illustrates the 6 parcellation levels of CHARM. The CHARM parcellation can be used for regional analyses of cortical morphometry at multiple spatial scales.

**Figure 7:**
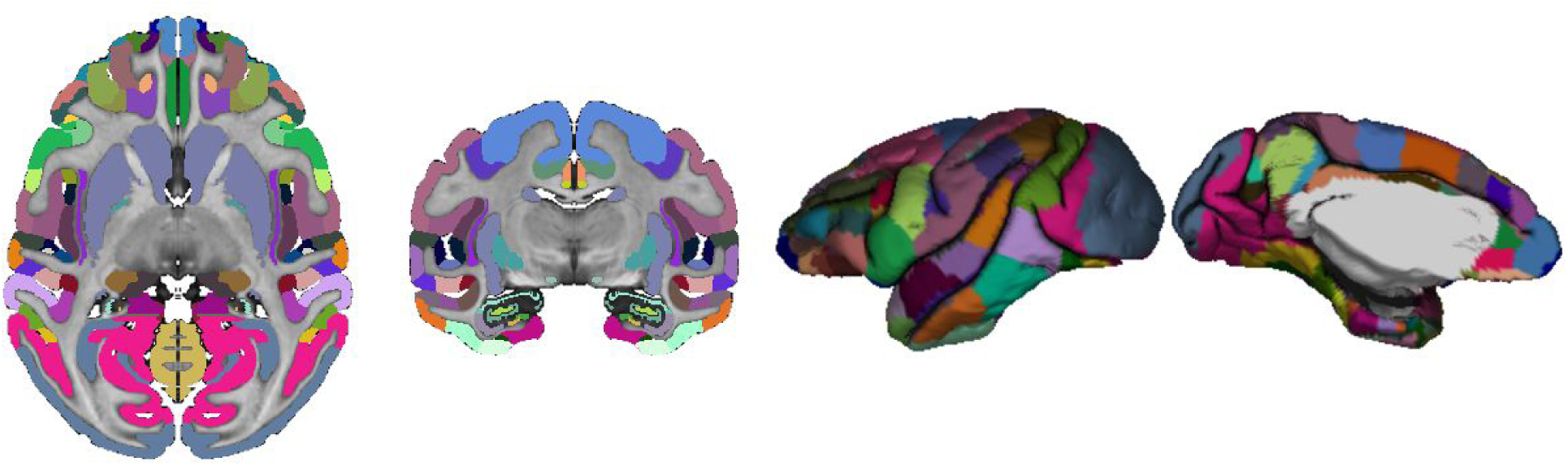
D99 volumetric segmentation defined on the anatomical scan of the D99 subject, transposed to the D99 registered pial surface.

**Figure 8:**
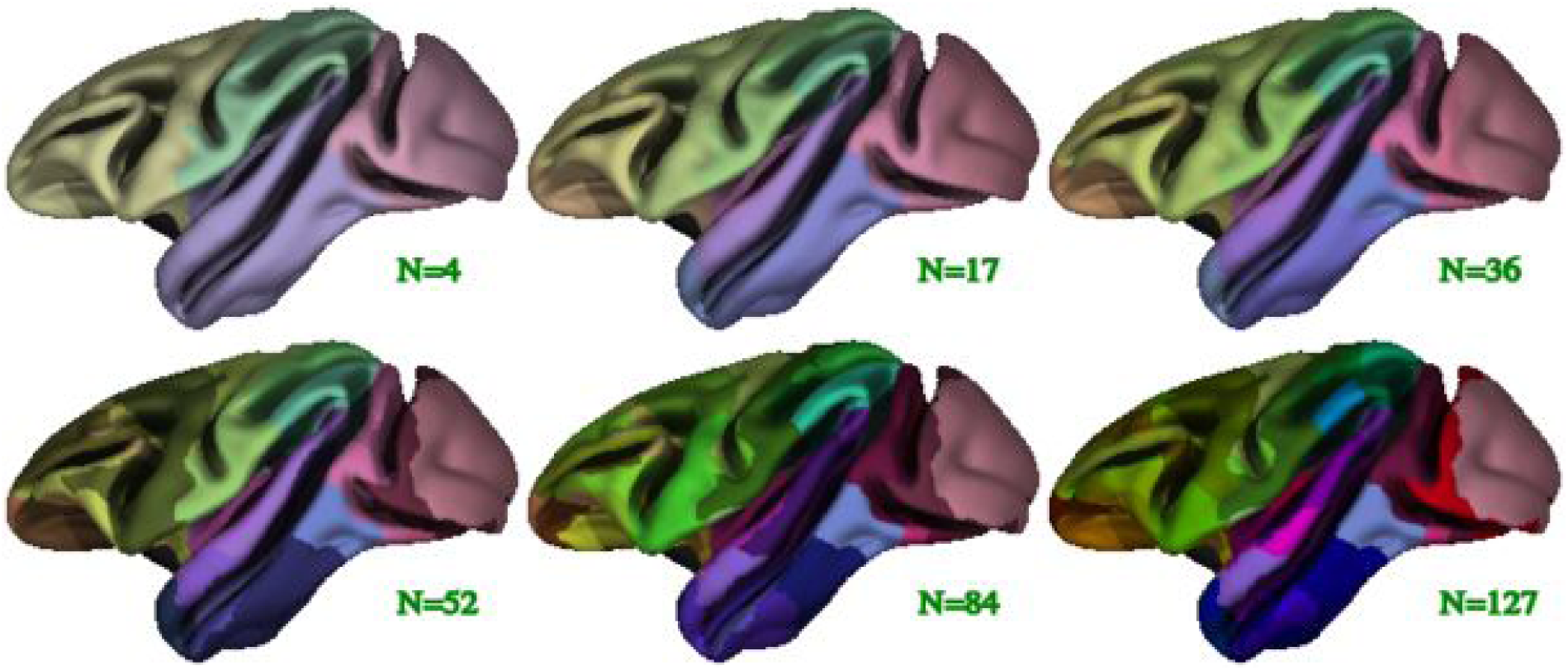
Illustration of the 6 levels of the symmetric CHARM parcellation viewed on the average symmetric mid-cortical surface for the NMT dataset. The CHARM parcellation is used in regional analyses by means of surface-based registration of individual subjects to the NMT surface template.

### 2.14 Quality control

CIVET produces a series of images to monitor the main processing stages of each subject. These images provide an efficient means for visual inspection of volumetric registration, tissue classification, surface extraction and convergence, surface-based registration, and cortical thickness. Moreover, several metrics are extracted (e.g. brain volume, mean cortical thickness, tissue percentages, etc.) and tabulated into a global csv file for detection of outliers within the population. Failed runs can be systematically identified at this level of quality control.

## 3. Results

### 3.1 CIVET preprocessing

The reliability of CIVET-macaque pipeline’s volumetric registration and brain-masking was tested on the T1w scans of the 31 macaques in the NMT cohort (Seidlitz et al. 2018) and 95 additional T1w scans of macaques obtained from the INDI PRIME-DE consortium (Milham et al. 2018). CIVET was run with the *without-surface* option, terminating after the stereotaxic registration, brain masking, and tissue classification stages. A value of 50 was used as the spline distance for N3 for all PRIME-DE scans, regardless of the field strength of the scanner, and a value of 75 for the NMT scans. A head height of 50mm, measured from the top of the head downwards, was used to crop the image to remove the neck for the purpose of registration.

Upon visual quality control, all brains were accurately registered to the NMT, without exception. The brain masking stage was successful for all subjects, except those from Site 10 for which the mask leaked into the scalp. For those 3 scans, the bone marrow inside the skull revealed signal intensities similar to GM and WM and was fused with cortex, without a visible inner bone border, causing the mask leakage. (The use of the T2w scan to correct the mask was preempted since the temporal lobes were truncated by the scanner field of view.) In a few other cases, from other sites, the mask touched the edge of the orbits or the bottom half of the cerebellum was truncated from the field of view of the native scan, but such small defects do not have any incidence on cortical surface extraction. Figure 9 reports the volume of the brain masks in standardized space. Agreement of the individual brain mask volumes with that of the NMT (red line) confirms successful registration to the NMT and brain extraction. The few outliers from site 10 are clearly visible in Figure 9. The mean stereotaxic brain mask volume for the PRIME-DE subjects, excluding the 3 outliers from site 10, is 92.9 ± 3.0cc, in agreement with the volume of the NMT brain mask of 92.8cc. Figure 10 shows the scan acquisition voxel size within and across sites. Scans for which all voxel dimensions were large (sites 12-14) generally had larger brain mask volumes and greater inter-subject variability. These failure rates are comparable to those observed in studies of human scans.

**Figure 9:**
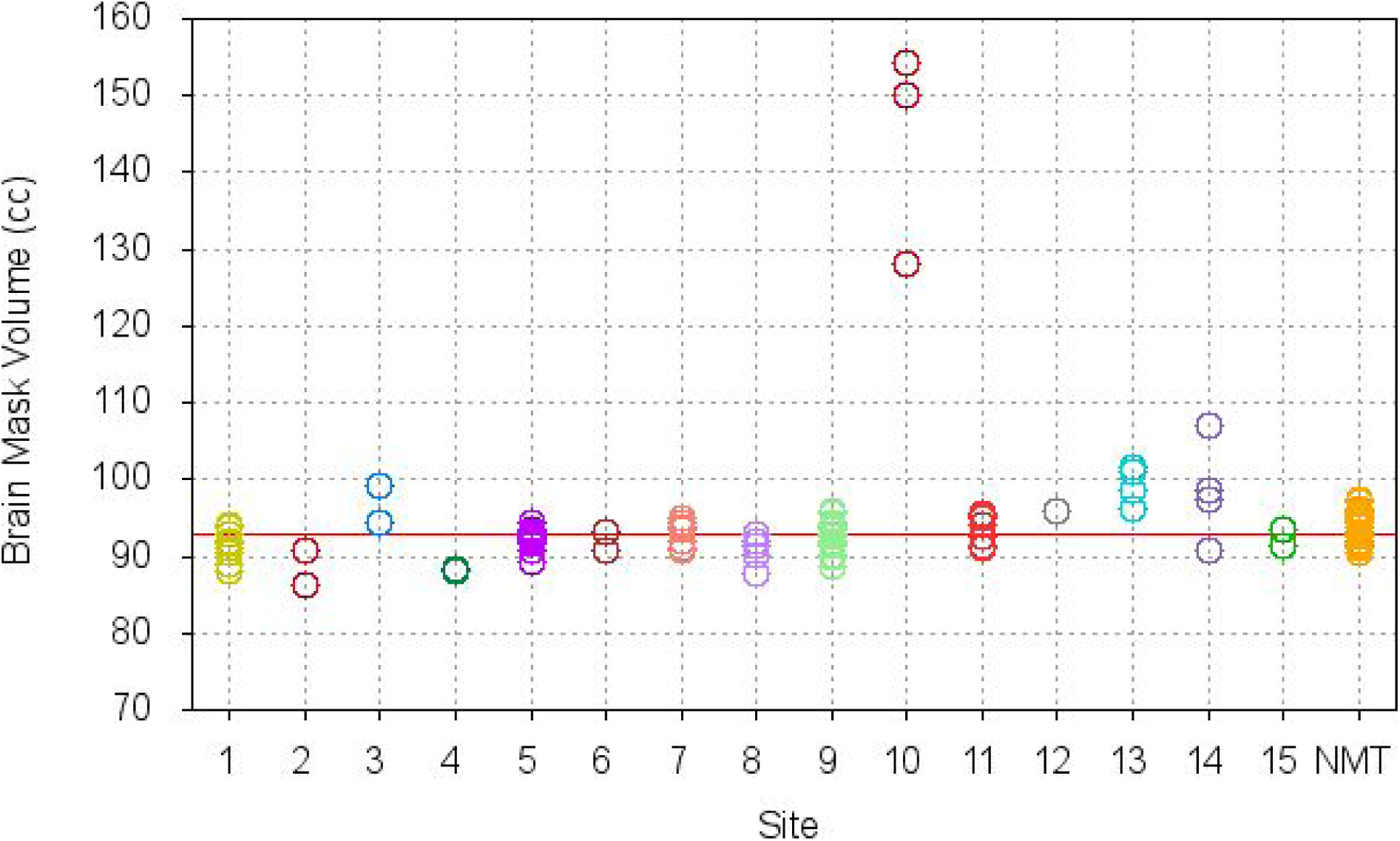
Brain mask volume in NMT standardized space for N=95 scans from the INDI PRIME-DE consortium and N=31 scans from the NMT cohort. The brain mask volume for the NMT volume is 92.8cc (red line). The small standard deviation for the volume of the brain mask in standardized space attests to the robustness of the brain extraction tool and the reliability of the volumetric registration to the NMT. The few obvious outliers show the processing failures.

**Figure 10:**
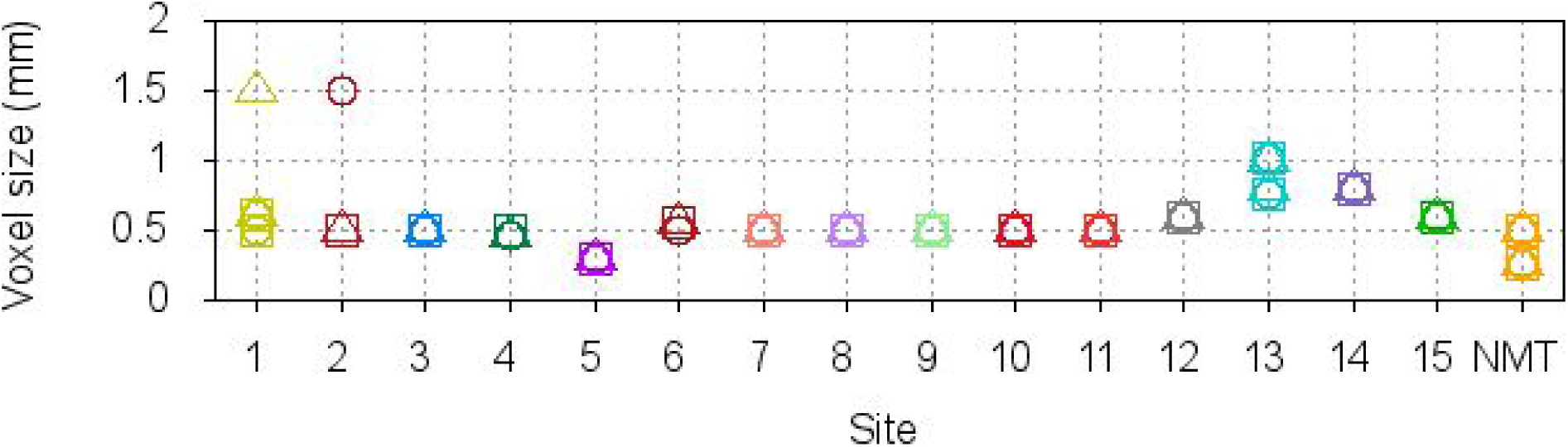
Acquisition voxel size for the N=95 scans from the INDI PRIME-DE consortium and N=31 scans from the NMT cohort. The three symbols (square, circle, triangle) refer to the voxel size in each of the 3 planes. By comparison to Figure 9, scans with a voxel size above 0.5mm tend to show higher deviation and variability from the volume of the brain mask of the NMT.

### 3.2 Cortical surface generation

Cortical surfaces were obtained for 29 of the 31 subjects in the NMT cohort (same two exclusions as for the creation of the average surface templates). The scans were processed at 0.25mm voxel size in stereotaxic space and the surfaces were produced at 40,962 vertices per hemisphere. A value of 75 was used as the spline parametric distance for N3. The *tlaplace* metric was used for the calculation of cortical thickness, blurred isotropically at a fwhm of 8mm. The disk usage per processed subject is about 325Mb (for short data type scans) to 550Mb (for float data type scans). (The T1w image input data type was preserved during processing.) Processing time ranged from 15-18 hours per subject. (For pre-processing only without surface extraction, the processing time is about 1.5 hours per subject.)

Figure 11 shows an overview of the CIVET stages and results for 4 NMT subjects. The results display the same level of accuracy for all subjects. Surface contours superimposed on the volume (top right panel for each subject) generally showed faithful correspondence to the gray and white borders. For some NMT subjects, the gray matter surface did not fully extend to the CSF on the orbital frontal cortex (OFC) surface, especially around the gyrus rectus (not shown), leading to artifactually smaller OFC thickness in these cases. These errors are driven by segmentation errors due to scanning artifacts present for some of the NMT subjects and may not be an issue for other datasets. In many cases, the WM surface along the insula was slightly internal to the WM/GM border, lying along the lateral surface of the putamen, especially dorsally where the claustrum is very thin (see Fig. 11). These shifts in the position of the WM surface are caused by a relaxation of the WM surface intensity gradient correction adjacent to the claustrum. Cortical thickness values for the insula are likely slightly inflated in these cases. The WM surface in this region was, however, smooth and did not exhibit the ‘claustrum invagination problem’ (Autio et al. 2020). Nor were there issues with segmentation in such problem areas as the ventral and anterior claustrum or the temporal stem.

**Figure 11:**
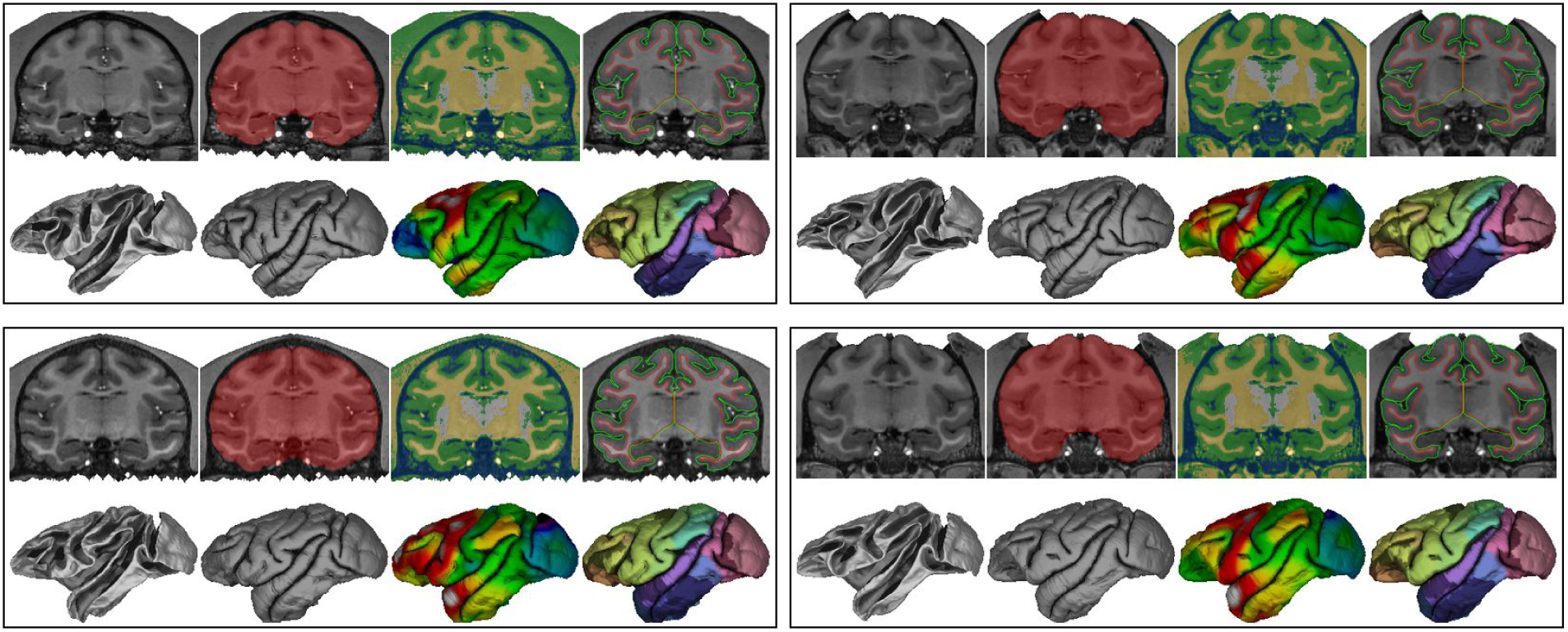
CIVET-macaque results for 4 NMT subjects. Within each set of images for a single subject, the top row, from left to right: coronal slice of the T1w image, brain mask, tissue classification (CSF in blue, cortical GM in green, portions of non-cortical GM in white, WM and other non-cortical GM in yellow), and cortical surfaces overlaid on the T1w image (WM surface in red, pial surface in green) bottom row, from left to right: white and pial surfaces; cortical thickness (range 1.0mm light blue to 3.0mm red) and the Level-4 CHARM parcellation viewed on the pial surface (52 ROIs).

Sulcal variability across the 4 subjects can be observed on both the WM and pial surfaces (Fig. 11, bottom row). Cortical thickness also exhibited some variability across subjects. The 3 subjects shown in Koo et al. (2012) show similar sulcal and thickness variability. In general, thickness increased from posterior to anterior.

#### 3.2.1 Cortical thickness

The average cortical thickness map across the 29 subjects and its coefficient of variation (COV=standard deviation/mean) are shown in Figure 12. The cortical thickness map patterns follow closely those obtained from human MRIs processed with CIVET, with cortical thickness increasing from posterior to anterior, as well as published macaque cortical thickness maps (Autio et al 2020; Calebrese et al. 2015; Donahue et al. 2018; Koo et al. 2012). The COV was low in most regions. Upon thorough examination of the cortical surfaces, the higher COV in some regions likely reflects a mixture of intersubject variability and limitations in the surface extraction process, namely the frontal pole (scanning artifacts due to nasal cavity), the temporal pole (presence of the middle cerebral artery), and the occipital lobe (lower gray-white contrast, owing to the high myelin content of this region [Donahue et al. 2018]).

**Figure 12:**
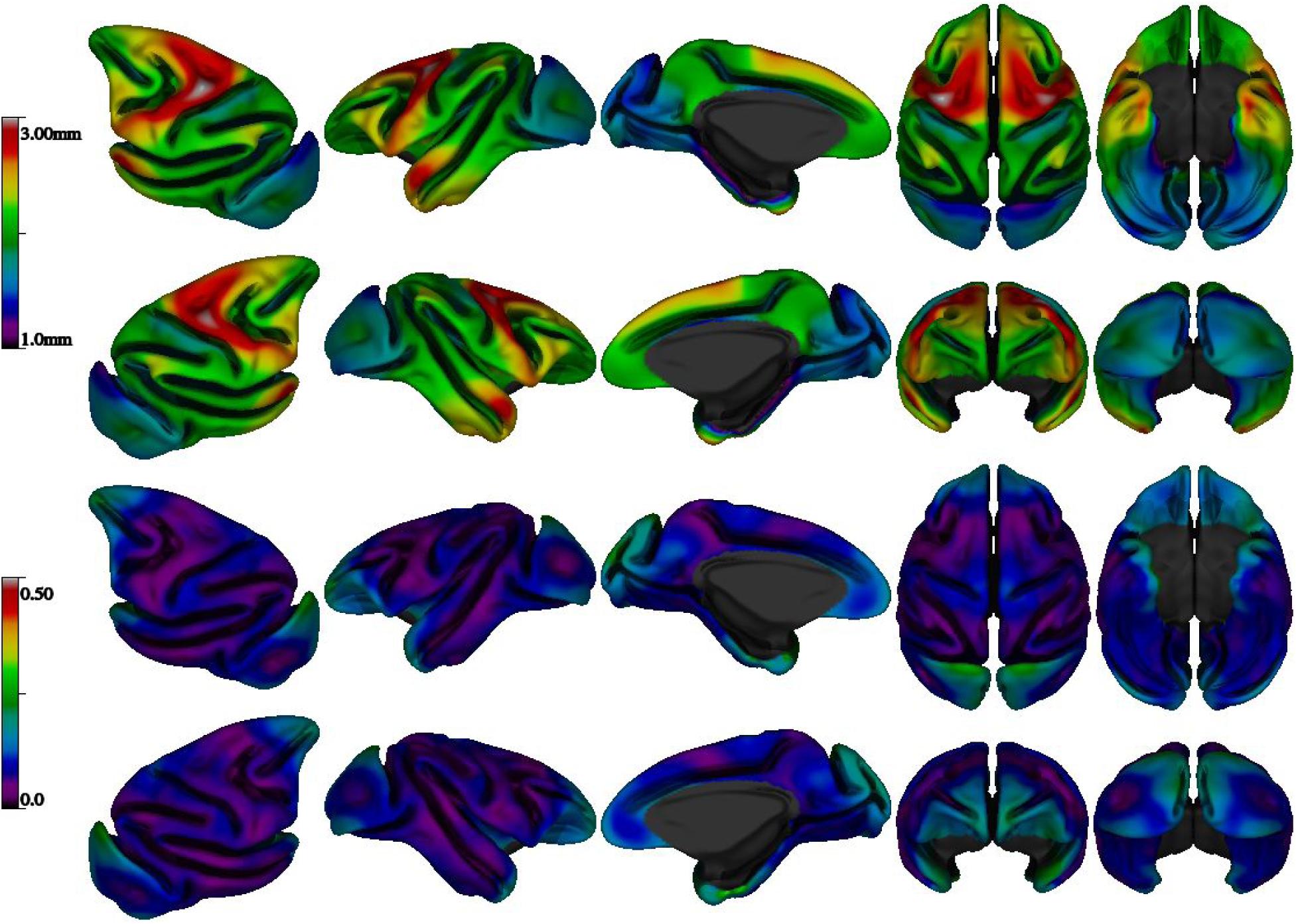
Mean cortical thickness (top) and coefficient of variation (COV) of cortical thickness (bottom) for N=29 subjects of the NMT dataset, using the tlaplace metric with fwhm of 8mm, viewed on the NMT average mid-cortical surface.

#### 3.2.2 Regional analysis

Regional cortical thicknesses based on the CHARM atlas are reported in Table 3. Thickness is reported for hierarchical atlas levels 1, 2, and 3 (with 4,17, and 36 regions, respectively) in the left and right hemispheres. Regional thickness was highly symmetric between the hemispheres. Mean cortical thickness, within the brain and averaged across the 29 subjects, was 1.99 ± 0.43 mm. The frontal lobe was thickest and most varied in its thickness across vertices (2.33 ± 0.43 mm), followed by the temporal (2.12 ± 0.41 mm), parietal (1.93 ± 0.31 mm), and occipital lobes (1.67 ± 0.25 mm). At level 2, particularly thick regions include the motor cortex (2.62 ± 0.36 mm), including primary motor cortex, premotor cortex, and the dorsal aspect of the cingulate gyrus, and the temporal pole (2.49 ± 0.49 mm). Particularly thin regions include the somatosensory cortex (1.89 ± 0.30 mm), the core and belt regions of the auditory cortex (1.79 ± 0.36 mm), and primary visual cortex (1.66 ± 0.22 mm), especially within the parieto-occipital sulcus and on the ventral surface in the inferior occipital sulcus. Figure 13 shows the histogram for the distribution of cortical thickness for several individuals and for the averaged cortical thickness across all 29 subjects (without blurring).

**Table 3:**
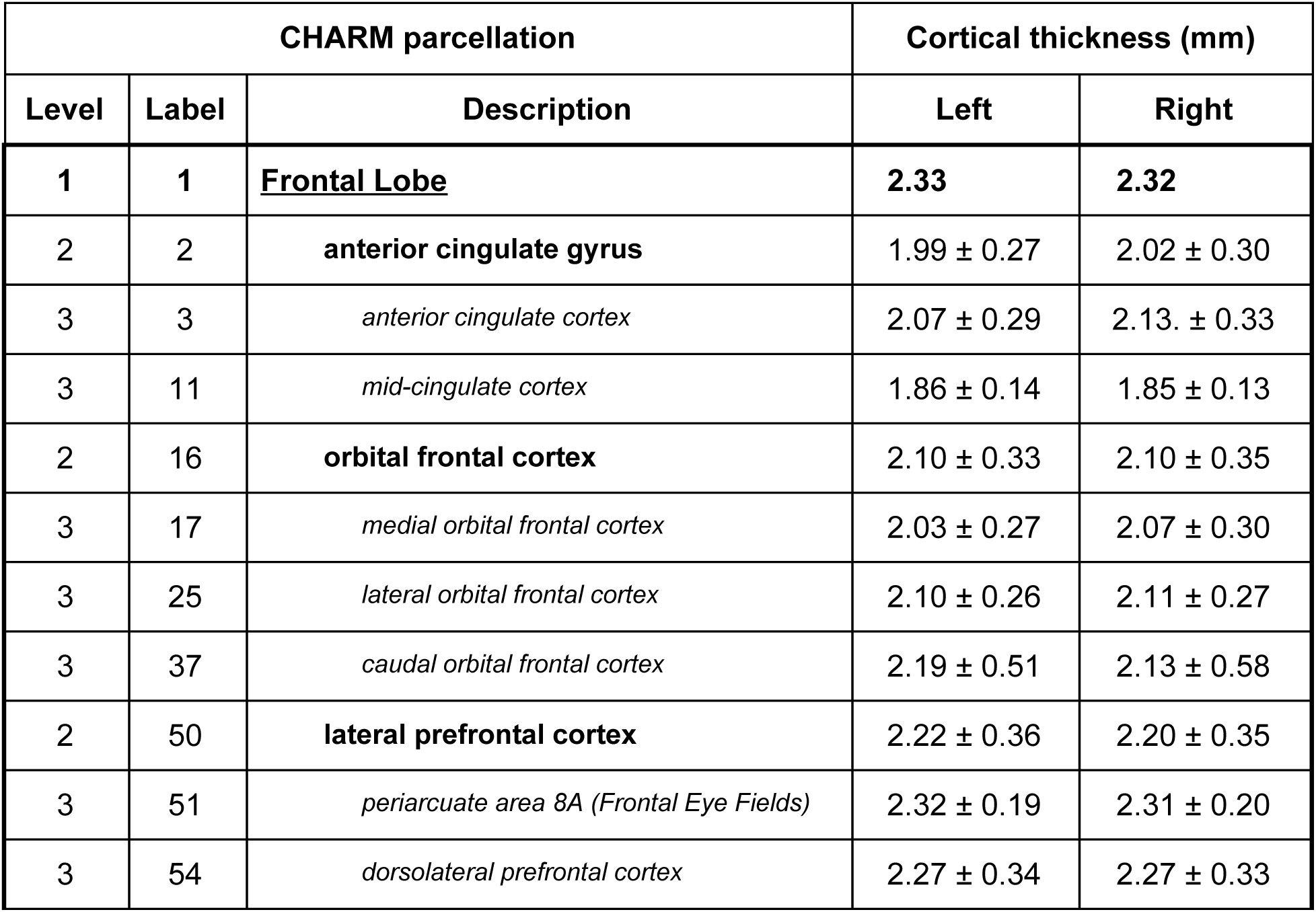
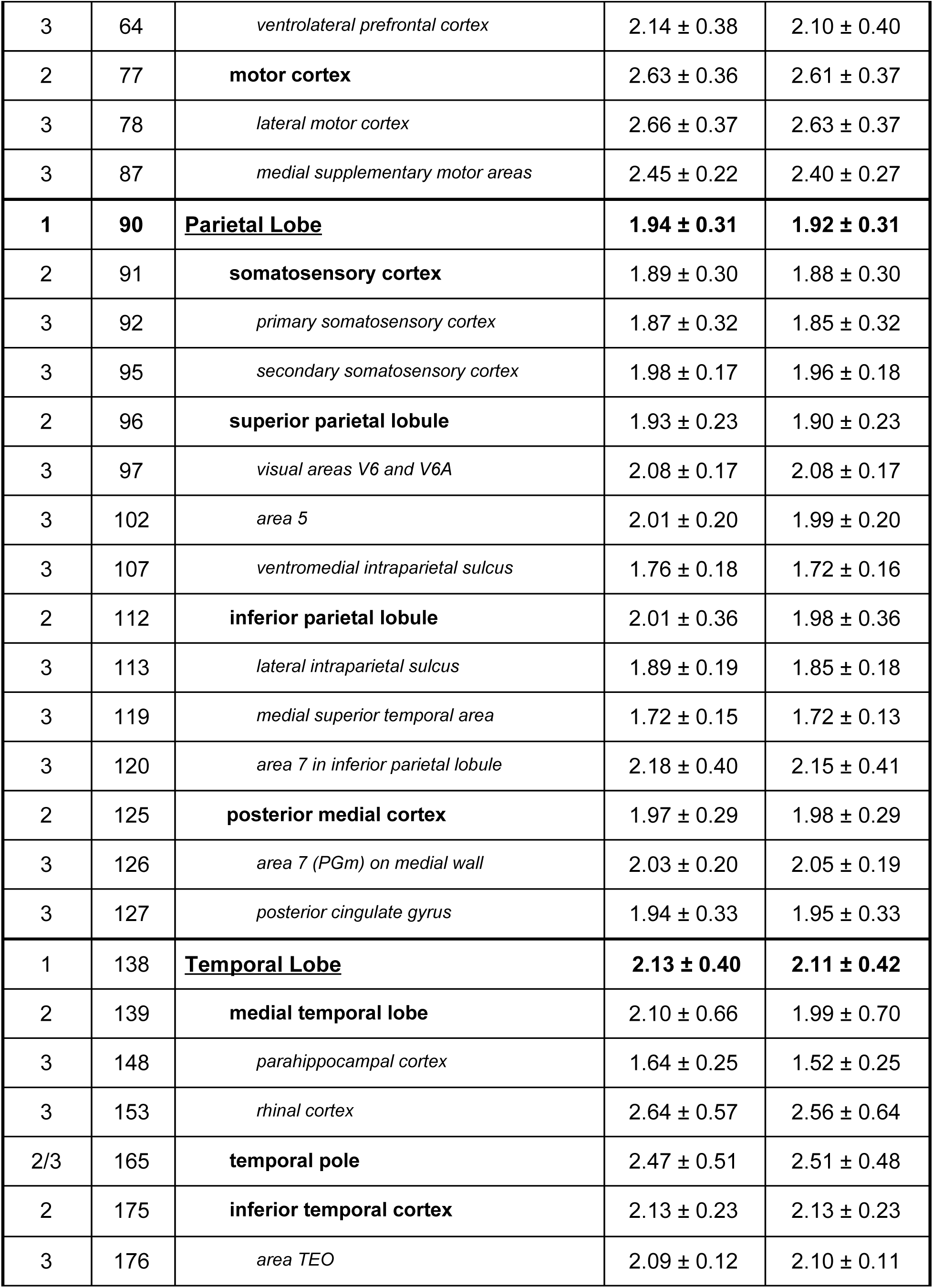
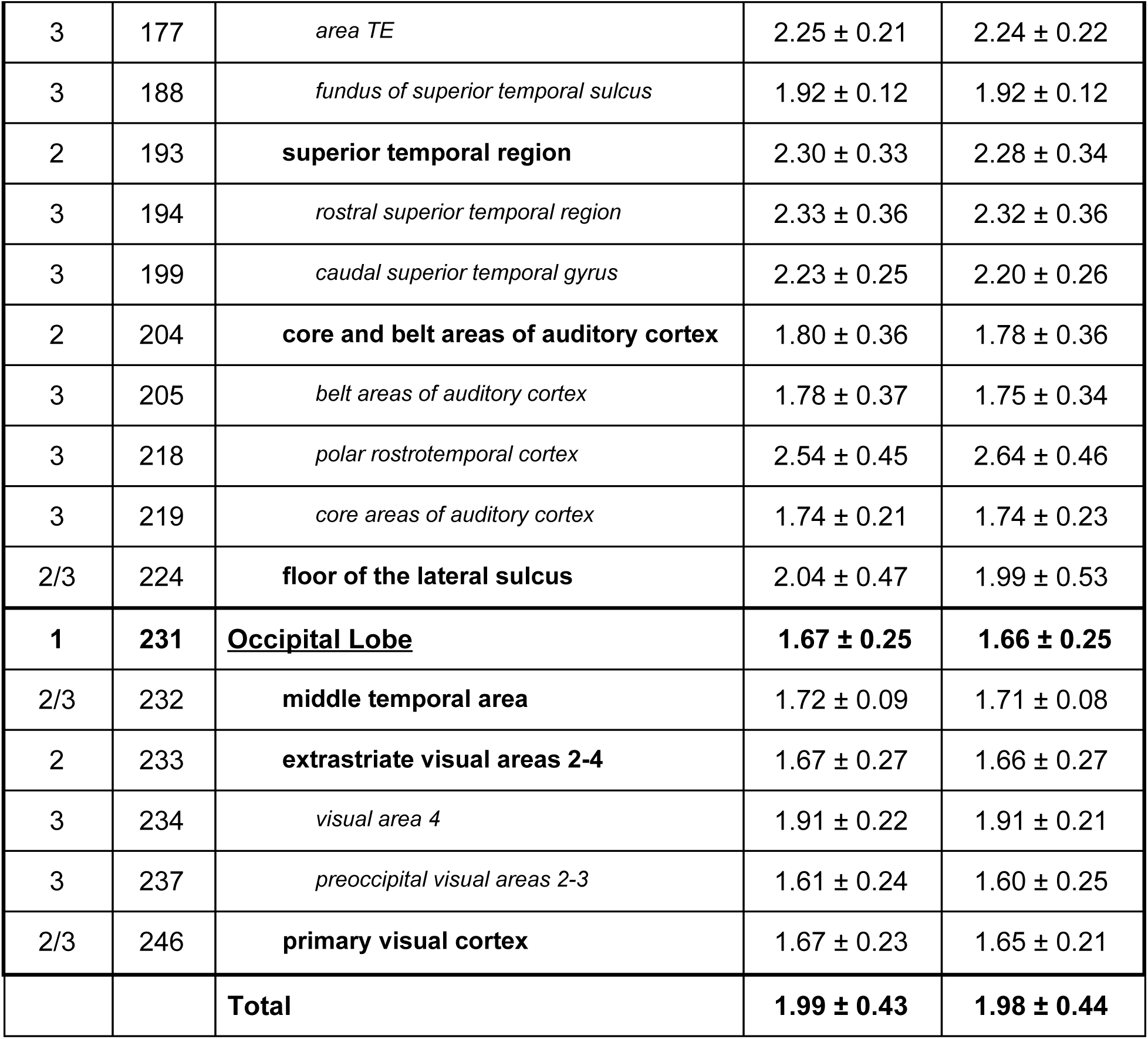
Regional mean cortical thickness of the NMT cohort (N=29) for the 3 broadest levels of the CHARM anatomical atlas, sorted by ordinal label number. Region names and atlas number labels are listed in different fonts and with different degrees of indentation depending on the CHARM level(s) in which they appear. Levels 1, 2, and 3, which contain 4,17, and 36 regions, respectively, are included. For each subject, the mean and standard deviation of the thickness (tlaplace metric with fwhm of 0mm) was computed across the vertices within each cortical region and hemisphere. Each region’s mean and standard deviation was then averaged across subjects. The overall standard deviation is higher than the standard deviation of individual specialized regions, indicating higher inter-regional variability than intra-regional variability. Thinner areas include somatosensory, auditory and visual cortices, while temporal pole and motor cortex were the thickest areas. The thickness of each region was nearly identical between the left and right hemispheres. The smaller cortical thickness values in the parahippocampal cortex (region 148) reflect its proximity to the hippocampus, which is not modelled. This region should be excluded in analyses.

**Figure 13:**
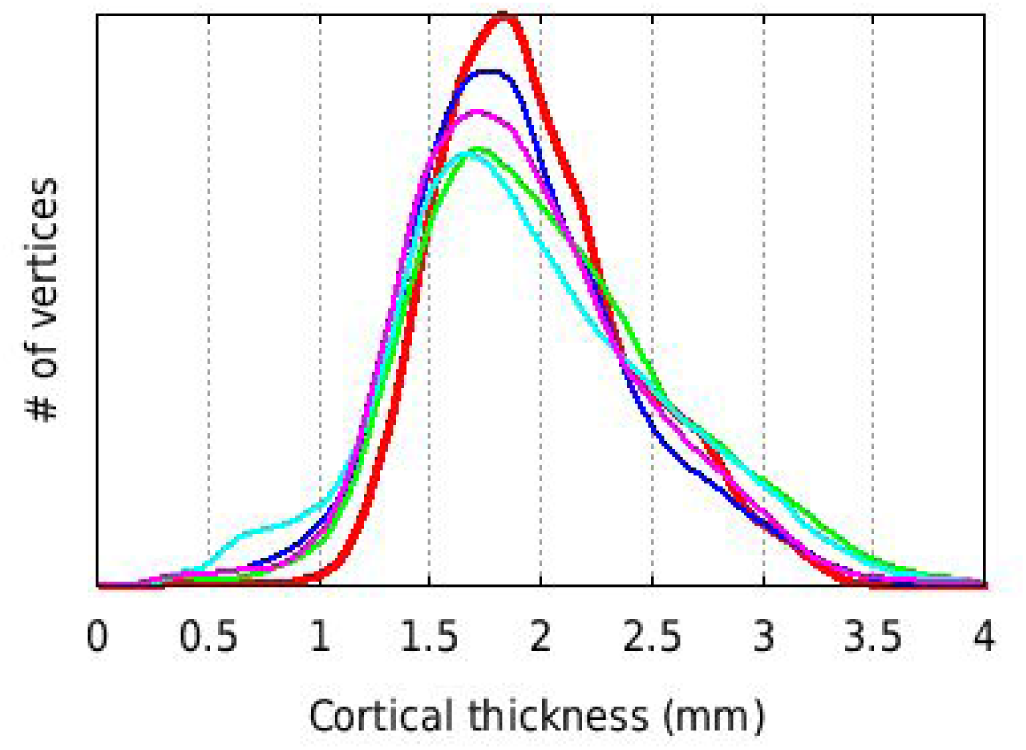
Histogram of mean cortical thickness for N=29 subjects of the NMT dataset (red thick line) and for 4 single NMT subjects (thin lines), using the tlaplace metric with no spatial smoothing (fwhm = 0mm). All 5 curves show the same total number of vertices (40,962 × 2 hemispheres) and area under the curve. The distribution for the mean curve is narrower (i.e., more central) than for most individual subjects. Consequently, the mean also has a higher peak than the other distributions.

#### 3.2.3 Surface area

The total area of the white matter and pial surfaces is displayed in Figure 14 for the NMT subjects (N=29). This total area includes the mid-plane wall cutting through the corpus callosum and the brainstem. Despite the inter-subject variability, areas of the left and right hemispheres are in close agreement within each subject for both cortical surfaces. The mean area per hemisphere was 10,098 mm^2^ for the white matter surface and 13,441 mm^2^ for the pial surface. These values are larger than those reported for the macaque (Autio et al 2020; Donahue et al. 2018; Koo et al. 2012), even when accounting for the mid-plane wall region. The larger values obtained here could be an indication of deeper sulcal penetration of the pial surface and increased morphological details captured by the extracted surfaces.

**Figure 14:**
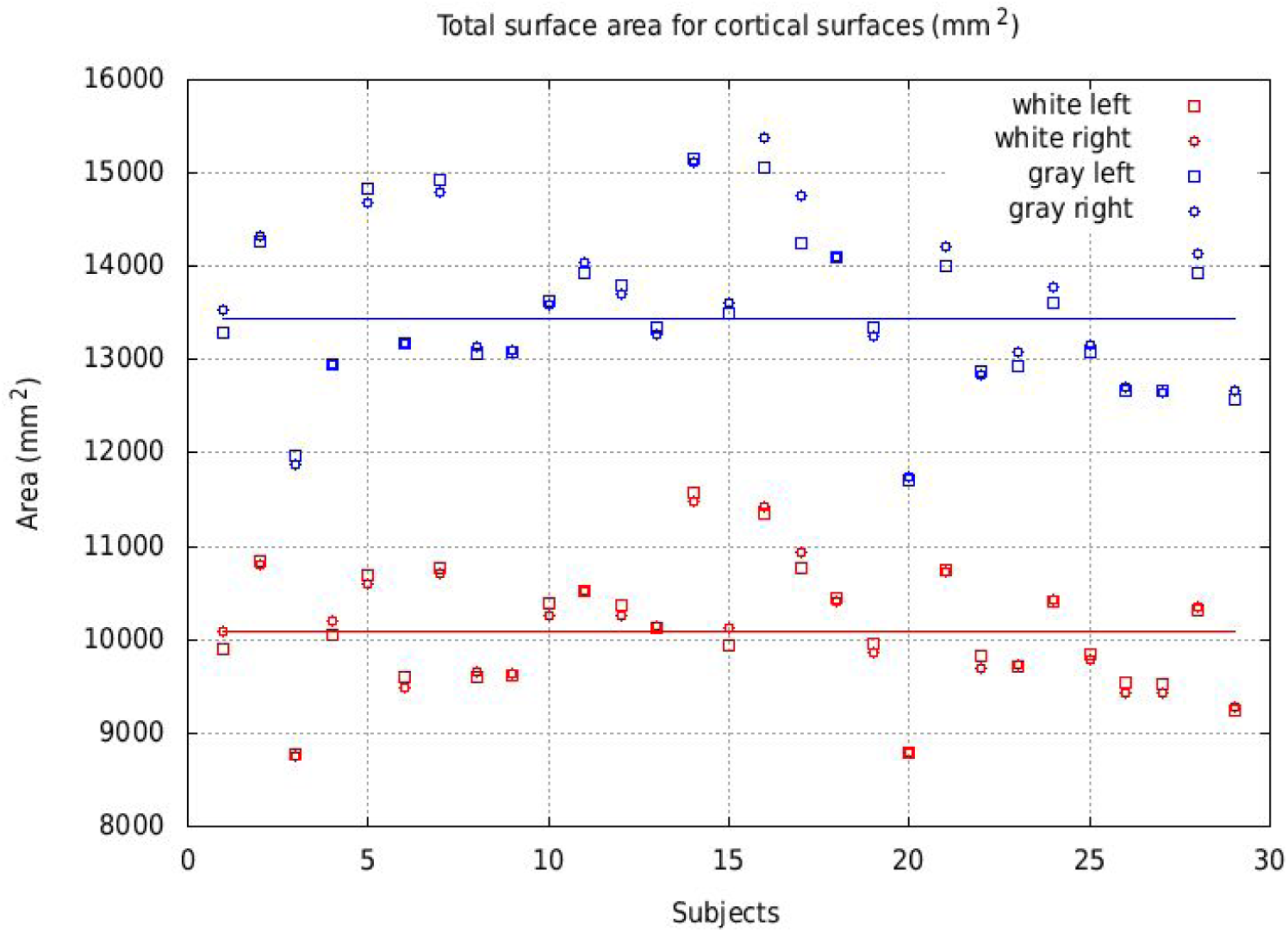
Total area for the white matter (red) and pial (blue) surfaces for the NMT cohort (N=29), for left and right hemispheres. The area per hemisphere, averaged across subjects and hemispheres, is 10,098 mm^2^ for the white matter surface (red line) and 13,441 mm^2^ for the pial surface (blue line). Left and right surface areas agreed for all subjects, independently of inter-subject variability.

## 4. Discussion

We have introduced CIVET-macaque, a pipeline for automated reconstruction of cortical surfaces of macaque brains. The pipeline requires only a T1-weighted MRI image to generate high resolution cortical surfaces, capable of intersubject surface-based coregistration with per-vertex correspondence between gray and white matter surfaces. Preprocessing and surface reconstruction were customized for macaques to address species related issues such as variations in head size and brain morphology.

The pipeline has been developed primarily on 29 scans out of the 31 subjects in the NMT cohort. Further tests were conducted on a larger heterogeneous collection of 95 scans from the INDI PRIME-DE consortium. The success rate depended, inevitably, on the quality of the imaging data. Input voxel resolution, field inhomogeneities, the presence of visible blood vessels, and scanning artifacts can all potentially cause various defects in the cortical surfaces. Supplementary Figure 1 shows the typical variability amongst scans and some common artifacts and their repercussion on the outcome of the pipeline. Some scans have a desirable high GM-WM contrast, but suffer from scanning artifacts anterior to the frontal and temporal poles. These artifacts interfered with surface placement, in particular for the NMT cohort, causing the COV for cortical thickness in these regions to peak. Some other scans were mostly free of scanning artifacts, but had low GM-WM contrast, which proved problematic during the extraction of the white matter surface (intensity gradient correction). Still other scans were of acceptable quality and produced expected results. It is hoped that scanning protocols better suited to corticometry analyses will be employed in the future for additional structural data acquisition in monkeys.

Overall, the pre-processing phases of CIVET-macaque proved to be highly reliable and robust with regard to registration and brain-masking, independently of the quality of the scans. Cortical surface extraction was also successful although more challenging, being very sensitive to the quality of the data and, in particular, to the presence of high intensities representing blood vessels and other acquisition artifacts. The availability of a T2w scan could be used, in the future, to improve the detection of such blood vessels. Finally, sufficient scan resolution (0.5mm voxel size or smaller) is necessary to delineate the fine structures of the macaque brain morphology, unlike the human brain for which a scan resolution at 1mm voxel size has been employed for many years.

At present, to circumvent possible shortcomings in the white matter surface extraction, a brain mask with special annotations can be supplied with the input T1w scan, in native space. This initial custom brain mask is obtained from a first run of the pipeline and annotated in problematic areas where errors occurred during the white matter surface extraction. Label=1 represents the underlying brain mask. Label=2 represents offending misclassified voxels that should not be white matter (e.g. blood vessels, meninges/dura, gray matter with high signal intensities). Label=3 represents missing white matter voxels (e.g. reconstructed thin parahippocampal gyrus or incompletely filled ventricles for the purpose of white matter surface extraction). Basically, the annotated mask allows construction of a valid white matter mask for the white matter surface extraction by retaining classified WM within the cerebrum (label >= 1), while removing voxels incorrectly labeled as WM (label = 2), and adding missing WM voxels to preserve the spherical topology of the cortical mantle (label = 3). Hollow ventricles can also be closed completely (label = 3) to avoid leakage of the white matter surface into the ventricles. CIVET is restarted with the new annotated custom mask and new surfaces are obtained.

All 31 NMT T1w scans revealed high-intensity voxels representing blood vessels or other scanning artifacts causing various white matter surface extraction errors of different degrees. An annotated mask, in native space, was thus needed for each of the 29 NMT scans included in this study. The task was fairly quick and amenable for this small dataset. After the masks were annotated, the pipeline was restarted in a fully automated way. The ensuing white matter surfaces were of high quality, as specifically needed for the generation of the average white matter and mid-cortical surfaces with minimal variability due to algorithmic noise.

The generated maps of cortical thickness for the NMT cohort (Figure 12 and Table 3) revealed expected morphological trends. These include thinner somatosensory, visual and auditory areas, thicker temporal pole and motor cortex, with the remaining homotypic isocortex exhibiting thicknesses in between these extremes. This pattern is consistent with existing studies of macaque MRI morphology (Wagstyl et al. 2015) and similar to patterns seen in human histological studies of cortical thickness in the human BigBrain (Amunts et al. 2013; Wagstyl et al. 2020). CIVET-macaque would be an essential step towards creating an equivalent 3D histological atlas of the macaque brain.

The thickness map in Fig. 12 is qualitatively similar to that of published maps of individual subjects (see Koo et al. 2012; Oguz et al. 2015; Fig. 4 of Autio et al. 2020) and group averages (see Fig. 3 of Koo et al. 2012, n=8; Fig. 4 of Autio et al. 2020, n=12; Fig. S2 of Seidlitz et al. 2017, n=31), in particular that in Fig. S4 of Donahue et al. (2018, n=19) and Fig. 6 of Calebrese et al. (2015, n=10). The mean cortical thickness (*tlaplace*, fwhm=8mm) reported here is 1.99 ± 0.17 mm. The mean thickness of the NMT cohort agrees with that reported by Donahue et al. (2018, 2.0 ± 0.1 mm, n=19) and Autio et al (2020, 2.1 ± 0.1 mm, n=4), but is smaller than the value reported by Koo et al. (2012, 2.37 ± 0.19 mm, n=18). The pipeline employed by Koo et al. is a variant from an older version of CIVET. Their surfaces included on the order of 5000 vertices and the *tlink* metric was used for cortical thickness, which leads to an overestimation due to mesh distortion during the expansion of the pial surface. Note that the standard deviation, here, is the vertex-wise standard deviation across the n subjects, relative to the local vertex-wise mean of cortical thickness for the n subjects, averaged across the number of vertices on the surface (by hemisphere). It reflects how much cortical thickness varies locally - at the vertex level - across the subjects, relative to the local mean.

The variance in cortical thickness across the NMT cohort was low in most places. Greater COV (see Fig. 12) in the frontal pole, superior part of the occipital lobe and temporal pole appears in part to be driven by poor scan quality and algorithmic shortcomings. The frontal pole suffers from scanning artifacts due to the nasal cavity. In those subjects with the thinnest occipital lobe, the gray-white contrast was poor in that region, causing the true gray-white intensity transition to be missed and the white matter surface to be displaced towards the pial boundary. Finally, blood vessels (namely, the middle cerebral artery) in the vicinity of the temporal pole appear to be the source of the defects in both the white and gray matter surfaces in that region. Overall, the regions of high COV in this cohort are conjectured to be driven by processing errors and to be a weak indicator of true biological variance. Caution should be used when reporting results of biological relevance in these regions.

## 5. Conclusions

CIVET-macaque is the first true fully automated pipeline for processing macaque brains for corticometric studies. The pipeline, based on the imaging minc data format, has been developed in PERL and C/C++ for Linux systems. The open-source project (https://github.com/aces/CIVET_Full_Project) is also found on the PRIME-Resource Exchange portal (https://prime-re.github.io/). Outputs are produced in minc (HDF5-based) and NIfTI/GIfTI formats.

This new software enables the field of NHP imaging to further explore questions of structure and function across large populations and datasets in a new way. It is anticipated that the open usage of CIVET-macaque will promote collaborative efforts in data collection, sharing, and automated analyses from which the non-human primate brain-image field will further advance (Autio et al. 2020). For example, corticometry would provide a natural and complementary insight on longitudinal developmental studies in the macaque monkey (Scott et al. 2016; Wang et al. 2018). Finally, it is our intent to share complete CIVET outputs for the INDI PRIME-DE on the PRIME-RE portal (https://prime-re.github.io/) (Messinger et al., *this issue)*.

CIVET has the potential to be expanded to other non-human primate species, with possible extensions to the marmoset and the chimpanzee, using repositories of previously collected and, if possible, publically available anatomical scans. The ability to process marmoset, macaque, chimpanzee, and human data with CIVET will allow for comparative studies of surface-based morphometry.

## Data accessibility and availability

Scans used for the development of CIVET-macaque are already publicly accessible. These include the D99 post-mortem brain and atlas (https://afni.nimh.nih.gov/Macaque), the NMT volume (version 1.2; https://afni.nimh.nih.gov/NMT), and single subject scans from the INDI PRIME-DE repository (http://fcon_1000.projects.nitrc.org/indi/indiPRIME.html/). CIVET-macaque will be versioned and added to the global open-source CIVET project (https://github.com/aces/CIVET_Full_Project). Binaries for common Linux platforms will be provided for convenience. CIVET-macaque, the NMT, the CHARM (Jung et al., *this issue*), and updates to these resources can be centrally accessed from the PRIME-Resource Exchange portal (https://prime-re.github.io/) (Messinger et al., *this issue)*.

## CRediT authorship contribution statement

**Claude Lepage:** Conceptualization, software, data processing, formal analysis, writing. **Konrad Wagstyl:** Conceptualization, writing. **Benjamin Jung:** Resources, writing, software. **Jakob Seidlitz**: Conceptualization, resources, writing. **Caleb Sponheim:** Conceptualization, resources, writing. **Leslie Ungerleider:** Supervision, funding acquisition. **Xindi Wang:** Software and data curating, testing. **Alan C. Evans**: Supervision, funding acquisition. **Adam Messinger:** Conceptualization, resources, investigation, writing, validation, visualization, project administration, supervision.

## Declaration of competing interest

The authors report no competing interest.

## Acknowledgments

This research was supported (in part) by the Intramural Research Program of the NIMH and includes the relevant Annual Report number in the following format (ZICMH002899). Parts of this work have received support from Healthy Brains for Healthy Lives. KW is supported by the Wellcome Trust (215901/Z/19/Z). The authors report no conflicts of interest.

---

**Supplementary Figure 1:**
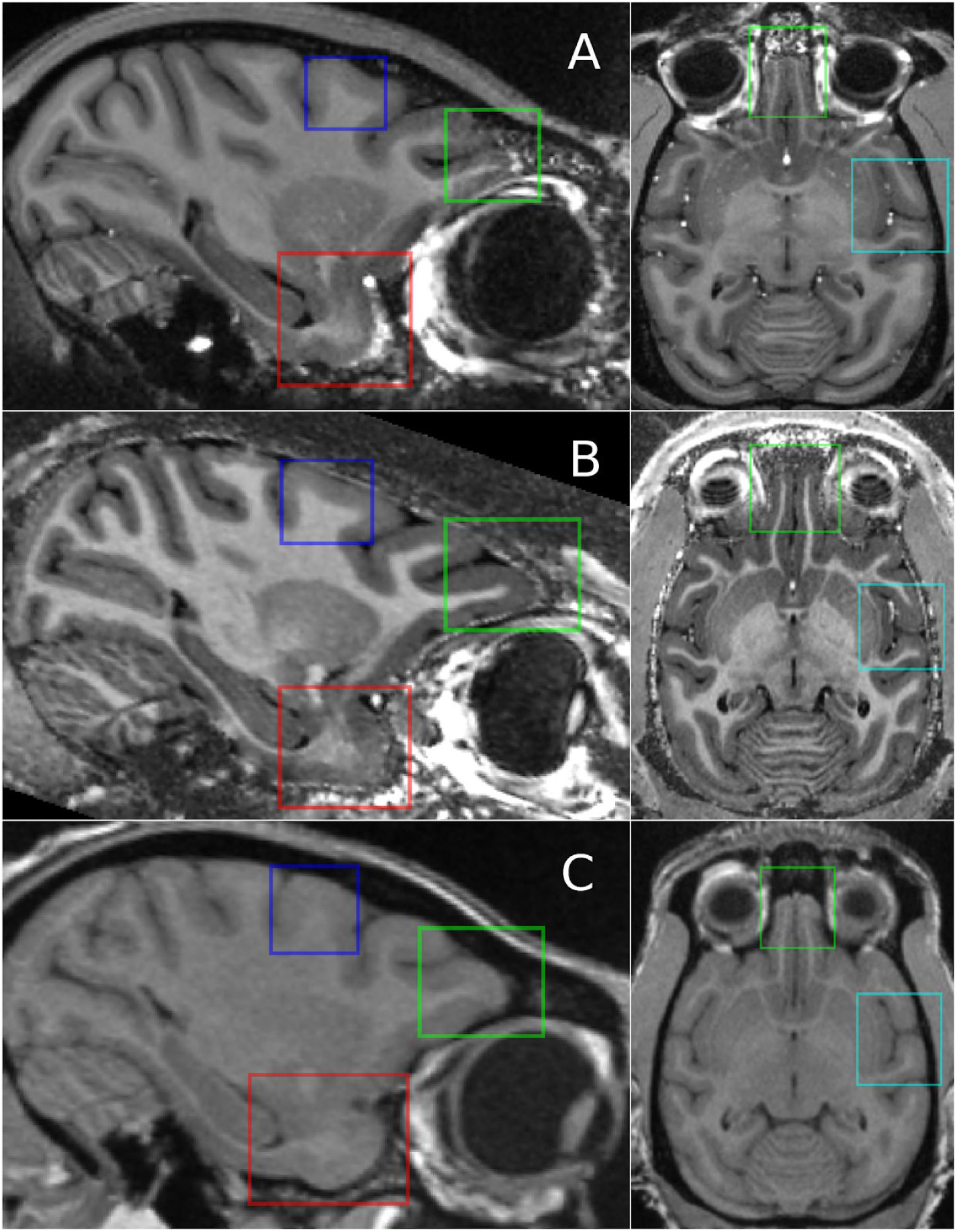
Scan variability and problem areas for 3 subjects from different sites (A: NMT subject, NIMH; B: PRIME-DE subject, site 10; C: PRIME-DE subject, site 11). GM-WM contrast (blue boxes) was clear for subjects A and B but very weak for subject C. Conversely, hyperintensities around the temporal pole (red boxes) and artifacts anterior to the frontal pole (green boxes), which likely arise due to proximity to the sinuses, were present for subjects A and B but not subject C. These hyperintensities diffused into the cortex causing local errors in the placement of the white matter surface. The cyan boxes reveal the presence of hyperintense voxels for blood vessels in subjects A and B (errors in white matter surface extraction), the weak GM-WM delineation of the claustrum in all subjects (misplaced white matter surface), and the presence of high intensity voxels for the bone marrow within the skull in subject B (leakage of brain mask into scalp).

## Notes

### Competing Interest Statement

The authors have declared no competing interest.

